# Predicting Organ-Specific Toxicity of Selective Androgen Receptor Modulators, using Transfer Learning on Graph Convolutional Networks

**DOI:** 10.1101/2025.08.27.672581

**Authors:** Alexander D. Kalian, Arthur C. Silva, Jaewook Lee, Jean-Lou C.M. Dorne, Claire Potter, Emilio Benfenati, Olivia J. Osborne, Miao Guo, Christer Hogstrand

## Abstract

Novel Quantitative Structure-Activity Relationship (QSAR) models were constructed using Graph Convolutional Networks (GCNs), to predict Drug-Induced Liver Injury (DILI), Drug-Induced Renal Injury (DIRI) and Drug-Induced Cardiotoxicity (DICT) of Selective Androgen Receptor Modulators (SARMs) – an emerging class of performance-enhancing drugs. Prior to training on DILI, DIRI and DICT datasets, the GCN QSAR models were first pre-trained on a variety of unrelated biomedical assay datasets, as an attempt to improve model performance via transfer learning. The success of the transfer learning was mixed; model performances were measurably improved via pre-training on certain datasets, by statistically weak increases. The optimal final QSAR models achieved overall accuracy scores of 68% for DILI (no significant improvement via ensemble modelling), 76% for DIRI (improved to 77% via ensemble modelling) and 65% for DICT (improved to 67% via ensemble modelling). Application of the most optimal singular models to a dataset of 25 SARMs predicted that 21 of the 25 SARMs are either DILI-positive, DIRI-positive, or both – which raises concern, given the rising use of SARMs. All SARMs except for one were predicted as DICT-negative. A novel definition of the Applicability Domain (AD) was used, intended for close relevance to the models, via generating three-dimensional graph embeddings, for each model. Convex hulls were fitted around training data embeddings, with a ±10% buffer, defining the AD as the region of embedded chemical space covered by the convex hull, for a given model. Subsequent analysis found that a majority of DILI, DIRI and DICT testing data lay within the AD, alongside a majority of the SARMs – adding consensus to the reliability of the predictions.

## 1 Introduction

### 1.1 Health Concerns of Selective Androgen Receptor Modulators

Muscle growth promoters and anabolic substances are being developed for the treatment of disease-induced loss of muscle mass and function [1]. The anabolic substances include Selective Androgen Receptor Modulators (SARMs) [2]. SARMs are an emerging class of performance enhancing drugs, which work by selectively binding to androgen receptors in certain tissues, promoting bone and muscle growth [2]. Given their selective nature, SARMs are intended as safer alternatives to anabolic steroids and other conventional performance enhancing drugs (which are known to have numerous adverse health effects) [2]. The adverse health effects of SARMs are however poorly understood, yet previous studies raise concerns around toxicological endpoints such as hepatotoxicity [3, 4], reproductive toxicity [5, 6], nephrotoxicity [7], and cardiotoxicity [8]. Despite these concerns, studies report that the use of SARMs is growing, recreationally through various supplements marketed to fitness enthusiasts [9] and reports of liver injury through their use [10]. Furthermore, there is emerging social media and SARMs abuse trends [11]. Google trends analytics for the term ‘SARMs’ suggest that SARMs’ search interest is at an all-time high [12].

In 2017, the FDA issued a public advisory stating that SARMs were being included in bodybuilding products and that these compounds posed an increased risk for heart attack, stroke, and liver damage [13].

Adverse outcomes, such as liver or kidney damage, emerge as the final stage of a toxicological Mode of Action (MOA), which consists of toxicokinetic and toxicodynamic processes, following an external dose [14, 15] (which in the case of SARMs, is typically via intravenous injection or oral consumption [16]). Please see Fig. 1 below:

**Figure 1:**
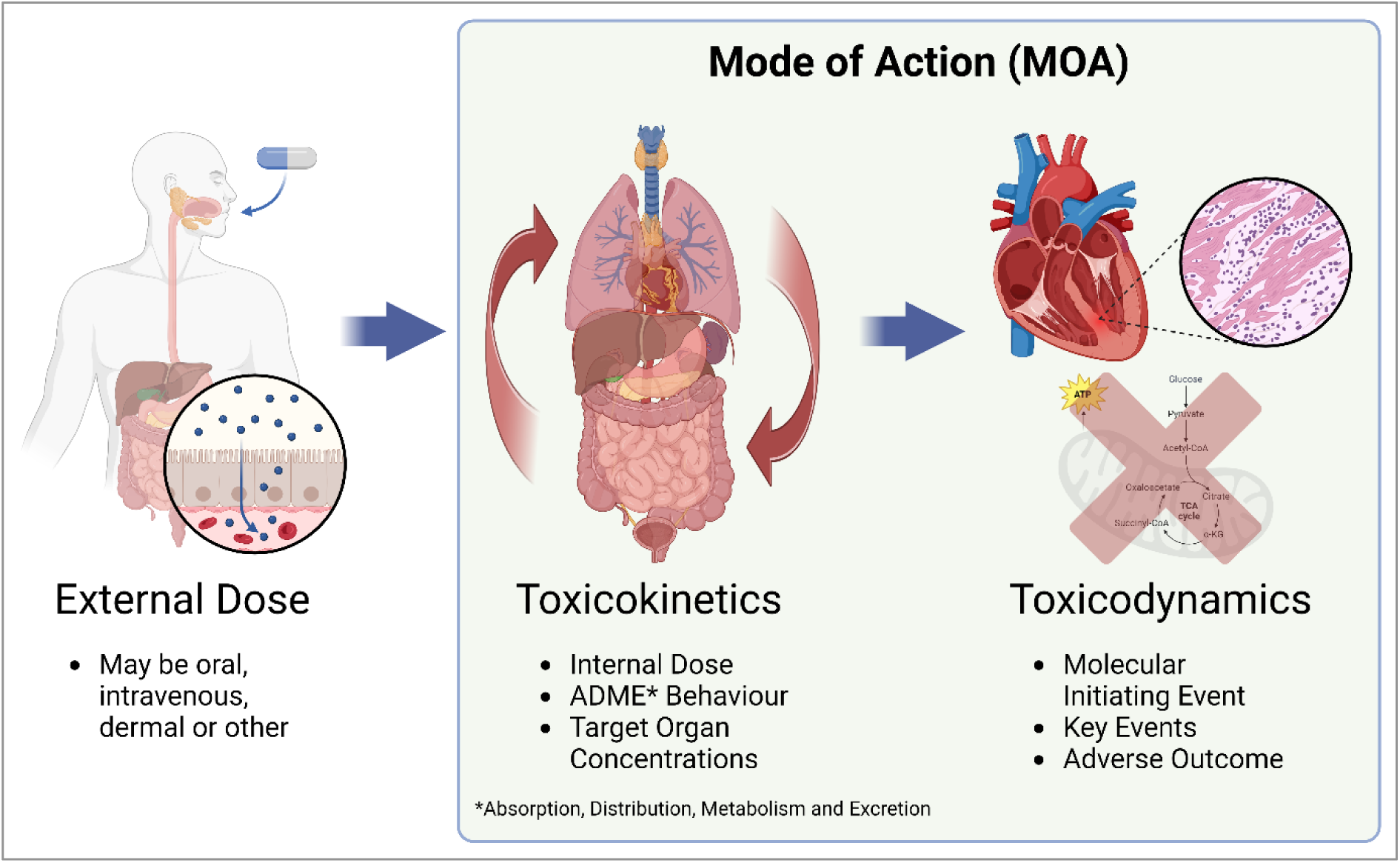
Overview of the different constituent stages of a toxicological MOA [14, 15]. Created in BioRender. Kalian, A. (2025) https://BioRender.com/sgl7srh

As per Fig. 1, toxicokinetic phenomena involves Absorption, Distribution, Metabolism and Excretion (ADME) behaviour of an internal dose of a chemical, resulting in target organ concentrations [14, 15]. Toxicodynamic phenomena subsequently concerns biochemical pathways on a cellular level, taking place within relevant organs and tissues [14, 15]. Toxicodynamic effects start with a Molecular Initiating Event (MIE) (often entailing competitive binding to an enzyme or receptor, although other types of MIEs also exist) and resulting in chains of biochemical reactions (or a lack of) which lead to an adverse outcome [17, 18]. In the case of a toxicological endpoint such as hepatotoxicity, specific adverse outcomes may include steatosis, cirrhosis or acute hepatic necrosis, among others [19].

### 1.2 Quantitative Structure-Activity Relationship Modelling using Deep Learning

Current chemical risk assessment frameworks are overly reliant on *in-vivo* animal studies [20], which pose numerous issues around ethics, validity, cost and future scalability. New Approach Methodologies (NAMs) attempt to mitigate these issues, including through *in-silico* approaches such as Quantitative Structure-Activity Relationship (QSAR) modelling [21]. QSAR modelling attempts to model biological activity of chemicals, as a function of molecular structure, while increasingly utilising Artificial Intelligence (AI) algorithms to learn relevant trends and relationships across toxicological space [21]. In particular, deep learning (typically via multi-layered artificial neural networks) has emerged as a leading form of AI in the cheminformatics space, utilised in a growing number of successful studies [21-23].

Molecules may be represented through a variety of different descriptors that AI algorithms may utilise, such as through Simplified Input Line Entry System (SMILES) strings of text (for language models) [24], as well as physicochemical descriptors [25], Molecular Access System (MACCS) keys [26], molecular graphs [27] and others [28]. Molecular graphs may be constructed to represent atoms as nodes, connected by bonds as edges, hence containing the full complexity of a molecular structure [27, 29]. Additionally, relevant physicochemical properties (e.g. atom electronegativity, bond order etc.) may be included as node and edge features [27, 29]. Molecular graphs are hence a powerful descriptor in cheminformatics, for encoding complex information about molecules, in a natural and intuitive form that closely aligns with the conventional representation of molecules as 2-dimensional graphs in the chemical and life sciences [27, 29]. However, conventional deep learning algorithms, such as multi-layer perceptrons or convolutional neural networks, require Euclidean-structured input data (usually in the form of a uniformly sized feature matrix) [30]; molecular graphs are however non-Euclidean, by nature [31].

### 1.3 Graph Convolutional Networks

Geometric deep learning is a branch of AI which encompasses deep learning architectures that are designed to handle non-Euclidean data [30]. Graph Neural Networks (GNNs) are an example of such, designed to directly operate on graph-structured data [30], including molecular graphs [27, 29, 31]. Graph Convolutional Networks (GCNs) are a popular GNN architecture [27], performing graph convolutions (analogous to convolutions performed by convolutional neural networks on image data and related) for message passing; this is used to build matrices of node features which are influenced by the node features of their regional neighbours in the graph, hence enabling the node feature matrix to acquire information indicative of the wider graph’s structure and the respective node’s position within that structure [32]. GCN message passing is defined via the following propagation rule between GCN layers [32]:

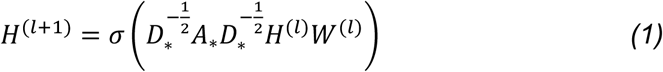

Where *H*^(*l*)^ is the node feature matrix for a given graph, at the *l*^*th*^ GCN layer (starting from an unmodified node feature matrix at *l* = 0). *σ* is an activation function (typically ReLU). 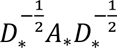 represents a normalised adjacency matrix for that graph, with self-looping for individual nodes, where *D*_∗,*ii*_ = ∑_*j*_ *A*_∗,*ij*_ (a diagonal degree matrix for the graph) and *A*_∗_ = *A* + *I*_*N*_ (the graph’s adjacency matrix *A* added to its identity matrix *I*_*N*_, to enable self-looping). *W*^(*l*)^ is a weights matrix for layer *l*, which is optimised via gradient descent.

The result of applying *k* number of GCN layers to a molecular graph, is a graph representation where the node features of each node (atom) contain information about other nodes up to *k*-hop degrees away [32] (see Fig. 2 below). The final node feature matrix may then typically be used to either make node-level predictions, or wider graph-level predictions, using feed-forward layers (i.e. a multi-layer perceptron). For graph-level predictions, it is commonplace to first perform pooling [27], to reduce dimensionality of the data into an efficient graph representation (types of pooling include mean pooling, maximum pooling and others) [33].

**Figure 2:**
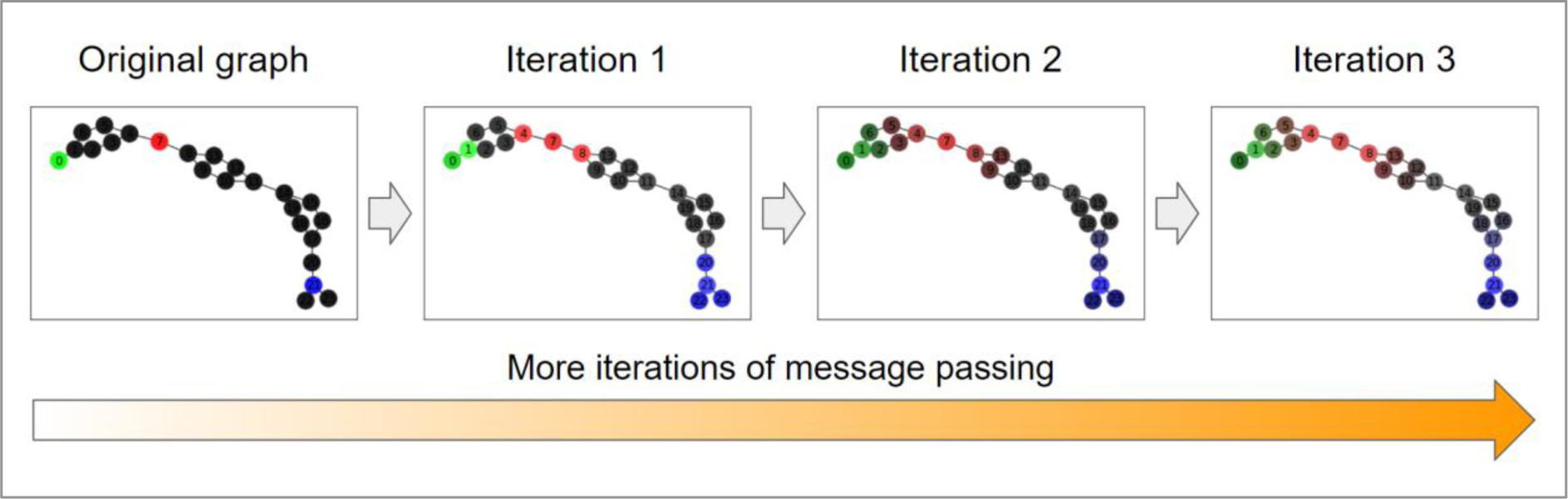
Visualisation of how message passing in a GCN (Equation 1) spreads node feature information to *k*-neighbours, over *k*-iterations, for node features of a molecular graph represented by RGB colour values. Note that, for this visualised example, no impact of the weight matrix is considered (which otherwise is trained to optimise the most useful spread).

### 1.4 Transfer Learning

Transfer learning is a technique used to improve the performance of AI models, especially in cases involving challenging or limited data [34] (as per Fig. 3 below). Transfer learning works by first pre-training an AI model on related tasks with abundant data, before then applying (either all or part of) the trained parameters to a model with a compatible architecture, as a starting point for subsequent training on the more challenging task [34]. Examples of previous studies that used transfer learning include a 2020 study which fine-tuned an encoder-based molecular structure prediction model to function as a QSAR model for predicting molecular activity [35]. An earlier 2018 study utilised transfer learning to create models for predicting pharmacokinetic parameters [36], whereas a 2021 study applied transfer learning to predictive QSAR modelling of compound reactivity to OH radicals [37].

**Figure 3:**
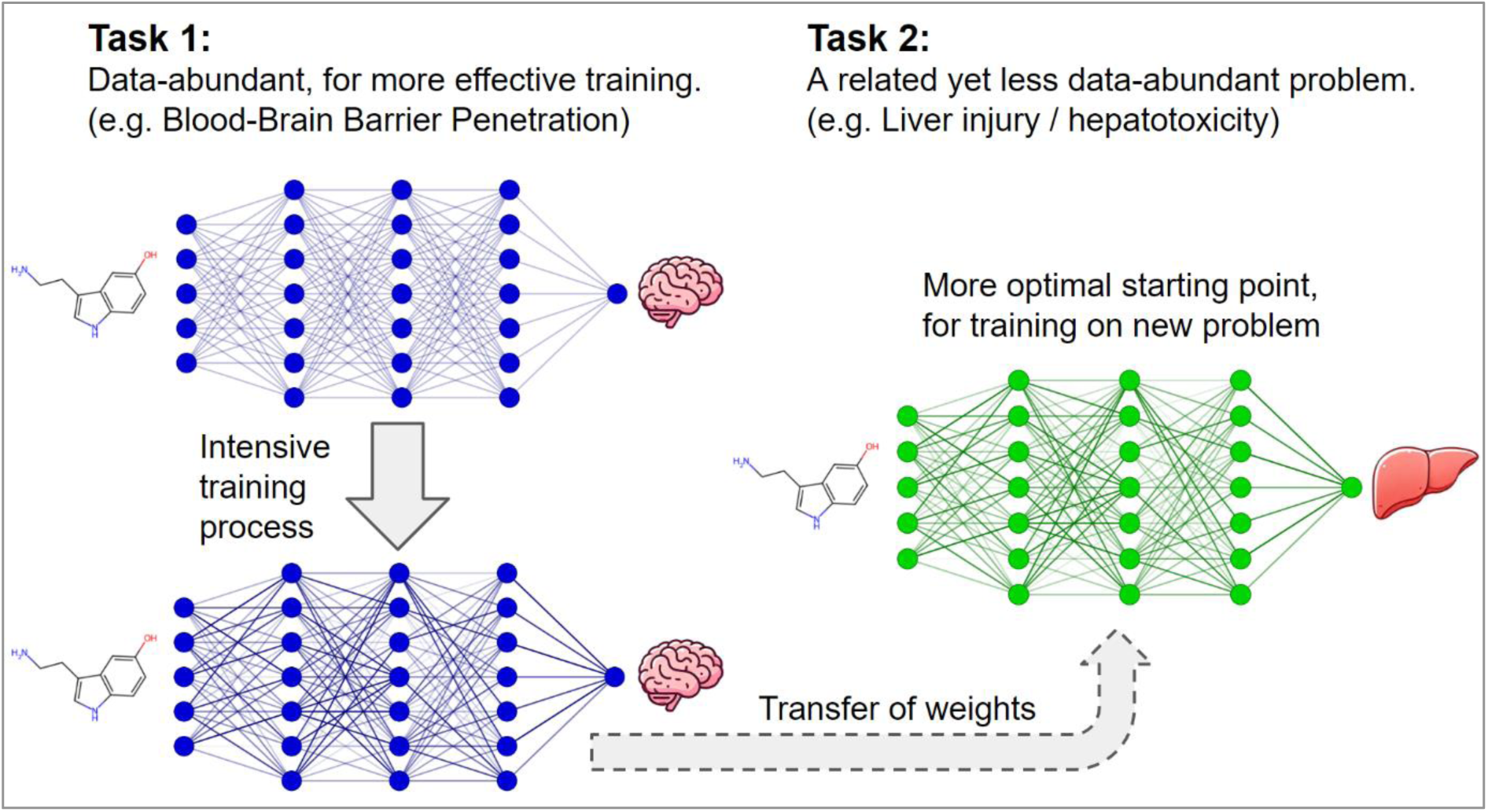
Schematic visualisation of transfer learning on deep neural networks, in the context of cheminformatics and predictive toxicology.

Transfer learning has also been specifically explored for GNNs, such as in a 2022 study (which explored use for GCNs and other GNN architectures) [38], as well as a 2021 study for highway forecasting [39], and a 2024 study which investigated transfer learning on GNNs for predicting bioaccumulation parameters of molecules [40]. A number of other studies, such as a 2020 study [41], a 2023 study [42] and a 2024 study [43] have however outlined that transfer learning on GNNs is often more fragile to failure than transfer learning over more established deep learning architectures (e.g. language models), with risks of negative transfer [42] - particularly when pre-training data [42], specific GNN implementation [43], means of fine-tuning [43], and other methodological considerations [41] are not carefully chosen.

Despite these challenges, transfer learning on GNNs, such as GCNs, using pre-training on generic biomedical data, may improve performance of toxicological QSAR models, even if potentially only marginally, for enabling organ-specific toxicity predictions of SARMs that are as accurate as possible.

The aim of this study is to construct such models, using transfer learning on GCNs, to output novel organ-specific toxicity predictions of SARMs, relevant to hepatotoxicity, nephrotoxicity and cardiotoxicity as toxicological endpoints.

## 2 Materials and Methods

### 2.1 Hypothesis Formulation

We hypothesise that use of GCNs will result in demonstrably effective QSAR models, across a range of different biomedical assay datasets, as well as organ-specific toxicity datasets. It is expected that transfer learning will lead to a measurable improvement in final model performance, even if potentially only minorly impactful. We furthermore predict that a sizeable number of SARMs will be predicted by these models as hepatotoxic and possibly also nephrotoxic and cardiotoxic, but that the predicted toxicity of the SARMs across these different endpoints may highly vary, owing to the inherently selective nature of androgen receptor binding by SARMs (and hence a highly selective occurrence of on-target adverse effects).

### 2.2 Data Collection and Pre-Processing

For the purposes of transfer learning, 9 biomedical assay datasets were obtained and curated. These datasets included: the benchmark Ames mutagenicity dataset by Hansen *et al.* (2009) (binary class-based) [44], the benchmark dataset for Blood-Brain Barrier Penetration (BBBP – binary class-based) included in MoleculeNet [45, 46], 2 randomly selected CompTox Chemicals Dashboard [47] assays (ATG_PXRE_CIS and CEETOX_H295R_OHPROG) which contained both Median Active Concentration (AC50) values and binary class-based active/inactive labels, as well as 3 randomly selected ChEMBL [48] binding assays for human proteins - Oestrogen Receptor (ESR1), Dopamine Receptor (DRD2) and Vascular Endothelial Growth Factor Receptor (VEGFR-2), respectively containing Median Effect Concentration (EC50), Inhibitory Constant (Ki) and percentage inhibition values.

Further to this, a dataset of 25 unique SARMs was obtained, via a systematic literature search [49-65].

Canonical SMILES descriptors for all associated datasets (which were lacking in various) were obtained via the online chemical database PubChem [66], via use of the PubChemPy Python API [67].

Organ-specific datasets were obtained. The Food and Drug Administration (FDA) Drug-Induced Liver Injury (DILI) dataset (DILIrank) [68] was obtained for hepatotoxicity, while a Drug-Induced Renal Injury (DIRI) dataset was obtained for nephrotoxicity, from the supplementary data of a 2024 study by Connor *et al.* [69]. Furthermore, a Drug-Induced Cardiotoxicity (DICT) dataset was obtained for cardiotoxicity, also from the FDA (DICTrank) [70]. Each of these datasets consist of classification-based data, for a variety of drugs. Although these endpoints are comparatively superficial, compared to the true complexity of toxicological MOAs and specific adverse outcomes for particular organs, the DILI, DIRI and DICT labels summarise entire toxicological MOAs, as they are based on clinical and experimental findings for organ-specific drug-induced injury in humans and animals (hence representing high-level overviews of whether or not each drug results in MOAs that significantly affect these organs).

Each of the organ-specific toxicity datasets contained molecules in canonical SMILES notation, as well as class-designation for each molecule, for a total of 4 different classes – these can in all cases be generalised as: high concern, slight concern, unknown and no concern. The data was binarised, for aiding in simplicity of the machine learning problem, by combining the “high concern” and “slight concern” data into a single “active” class (representing all molecules known to cause damage to the organ in question), whereas “unknown” data was excluded and “no concern” data was assigned to an “inactive” class. Similar approaches to binarising multi-class data have been carried out in other QSAR modelling studies [71, 72], including in our previous research [73].

Following this, basic curation measures were applied to all datasets, removing duplicate datapoints, as well as erroneous SMILES descriptors. Although all molecules were provided in canonical SMILES notation, consistency in formatting was enforced via standardising the SMILES, using the MolVS default ‘*standardize_*smiles’ algorithm [74]. Furthermore, all data split into 5 different folds, for later use in 5-fold cross validation, with class-balance in folds enforced through stratified random sampling of each class (with excess remaining data from a more abundant class discarded). 5 of the 9 biomedical assay datasets contained continuous data (rather than class-based data), hence in these cases stratified sampling for fold assignment took place by first organising the data in ascending order, then splitting the organised data into bins of 5 datapoints in size; random sampling was then performed over each bin, to assign a balanced representation of the *y*-data distribution to each fold.

Molecular graphs were constructed for all molecules, across all datasets. Molecular graphs were generated by first using the Python library pysmiles [75] to convert SMILES strings into graph objects compatible with the widely used library NetworkX [76], with subsequent processing taking place via use of NetworkX. The final molecular graphs contained atoms as nodes, with bonds as edges connecting the nodes, as well as node features representing physicochemical properties for each constituent atom; node features included atomic number, atomic mass, electronegativity, atomic radius, dispersion coefficient, dipole polarisability, fusion heat, proton affinity and number of implicitly bonded hydrogen atoms. Although no edge features were included, inclusion of the number of implicitly bonded hydrogen atoms as a node feature, for each atom in the molecular graph, was designed to de-facto hold information about the nature of the bonds formed with neighbours (discernible when considering valency rules for each element). Following construction of molecular graphs as NetworkX graph objects, all where subsequently converted to Data objects (which make use of efficient tensors) compatible with the GNN Python library PyTorch Geometric (PyG) [77], which also included associated, which also included associated *y*-data. In cases involving datasets which contained continuous *y*-data, the *y*-values were (unless already normally distributed) log-transformed (to enforce normal distributions).

### 2.3 GCN QSAR Model Overview

All QSAR models used identical architectures (see Fig. 4), for the purpose of enabling direct transfer learning between models. In all cases, 3 GCN layers were used, each containing 120 hidden channels, implemented via the PyG library, followed by global mean pooling (to aggregate node-level data into graph-level representations, for graph-level predictions). Following this, 3 fully-connected layers would make the final prediction, implemented via the PyTorch library, with the first two fully-connected layers containing 300 artificial neurons and the last layer containing just 1 artificial neuron. In the case of binary classification, the final output would represent the predicted class of either 0 or 1 (with the final prediction obtained by rounding up or down the output number). In the case of regression, the final output would represent the (scaled) *y*-value. The loss function used varied between classification and regression tasks; binary cross-entropy loss was used for classification models, whereas mean squared error was used for regression models. In all cases, Adam optimisation was used for gradient descent [78]. An overview of the GCN QSAR modelling architecture may be found below:

**Figure 4:**
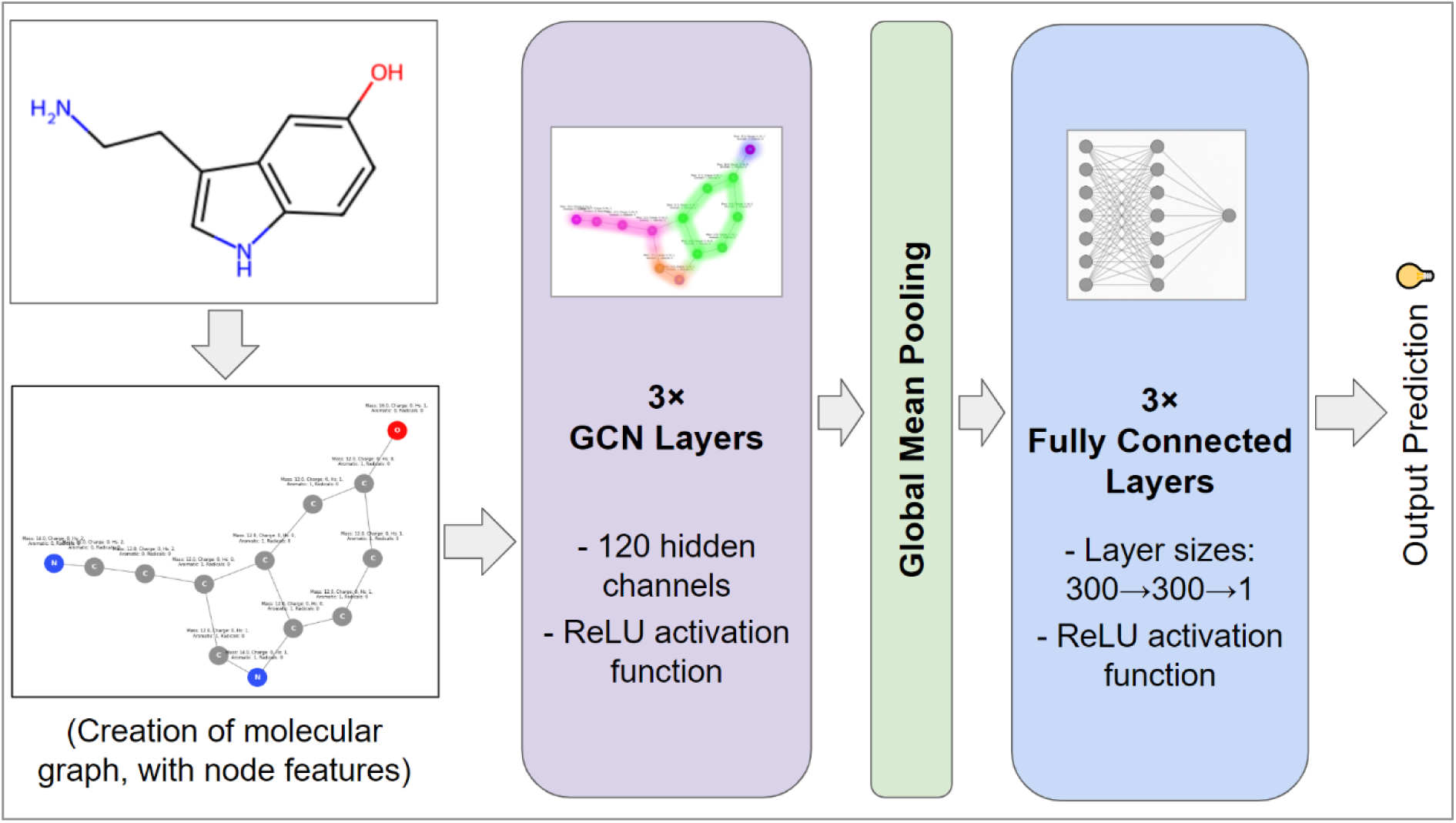
Overview of the controlled GCN QSAR model architecture used in this study.

All models were trained over 1,000 epochs. During the training process, the trained parameters of a given model would be saved as a local *.pth* file, with the file updated with new weights whenever an improvement to the loss was reached – hence the final *.pth* file would represent the most optimal trained state of that model.

### 2.4 Implementation of Transfer Learning

The GCN QSAR model architecture was first applied to training and testing on each of the 9 biomedical assay datasets, using 1,000 training epochs, repeated over 5-fold cross validation (with 1 fold used as testing data, in each instance). Once complete, the most optimal trained parameters for each biomedical assay dataset (using the most optimal fold-specific configuration) were then used as a starting basis for training and testing QSAR models of DILI, DIRI and DICT. Furthermore, as a control, subsequent QSAR models of DILI, DIRI and DICT were trained and tested without any prior pre-training.

### 2.5 Definition of the Applicability Domain

As per the OECD guidelines on QSAR modelling [79-81], all QSAR models must define the Applicability Domain (AD) - the region of chemical space which the model is applicable to, for new predictions. The guidance on how the AD should be defined is subject to broad interpretation, as demonstrated by broad variations in its definition across different studies; quantifying chemical similarity varies (e.g. physicochemical descriptors, MACCS keys etc.), while also considerations of the boundaries of the AD vary (e.g. hard boundaries around training data, density-based boundaries etc.).

For the purposes of this study, a novel definition of the AD is explored (Fig. 5), intended as naturally more representative of the GCN QSAR model used. Following training and testing of the GCN QSAR models, each optimally trained model would then be used to generate 120-dimensional molecular graph embeddings, using the trained GCN and global mean pooling layers. These 120-dimensional embeddings would then be reduced to 3-dimensional space (to aid in intuitive visualisation) using Principal Component Analysis (PCA). The boundaries defined by a convex hull, fitted around the 3-dimensional training data embeddings, for a given QSAR model, furthermore with a ±10% tolerance at the boundaries (in terms of distance of a convex hull vertex from the centre of the convex hull) would then be used to define the region of chemical space within the AD Although this definition of the AD requires available knowledge of the trained parameters of a given GCN QSAR model, it is reproducible as long as the trained models are provided in open-source (in terms of Python code and associated *.pth* files). A visualisation of this definition of the applicability domain may be found below:

**Figure 5:**
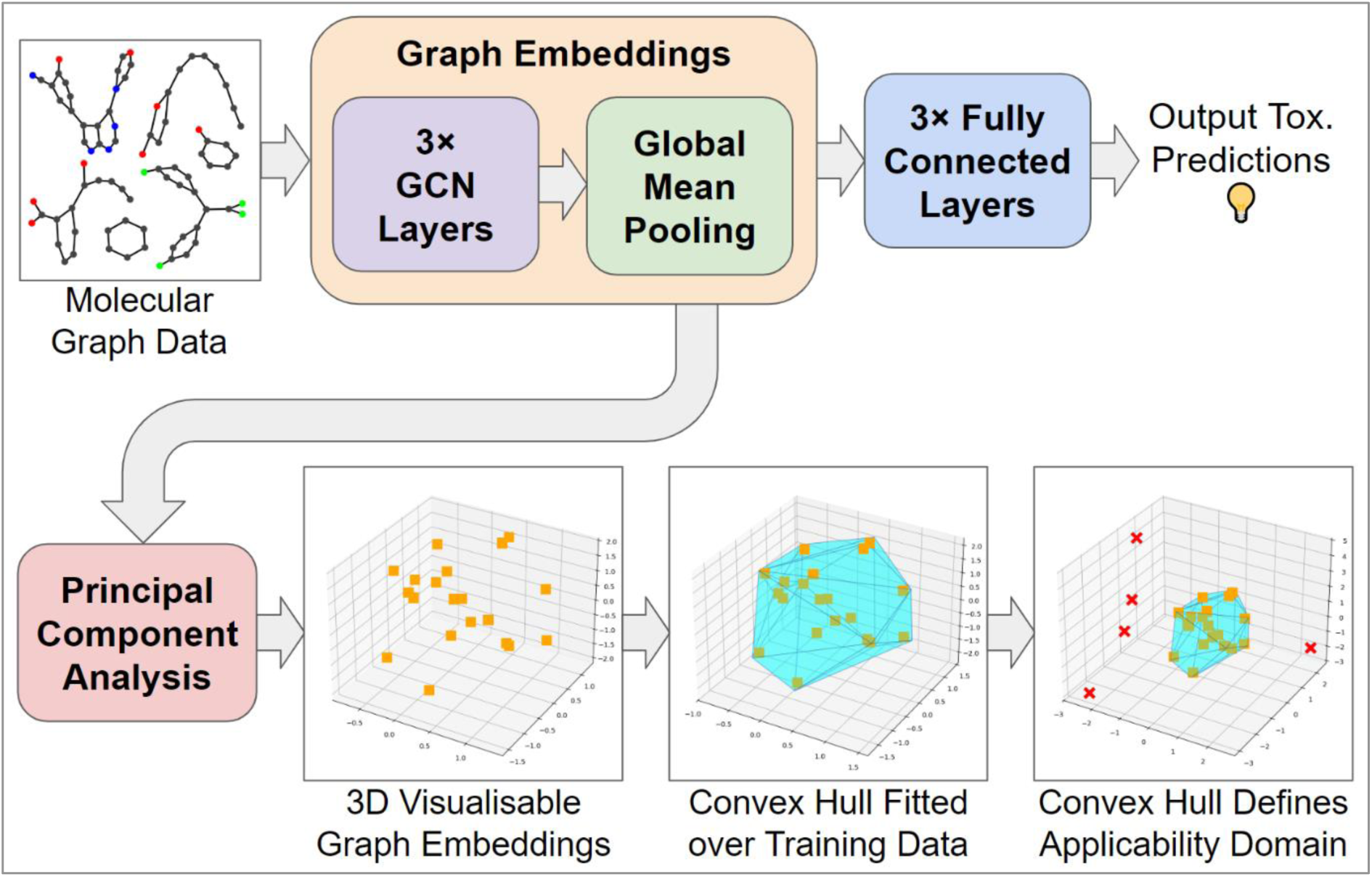
Schematic overview of how the AD is defined, in relation to the GCN QSAR model.

### 2.6 Exploration of Ensemble Models

Simple ensemble models were explored, via Ensemble Voting Classifiers (EVCs), combining the predictive power of all models of controlled endpoint and fold-configuration, but subjected to differing pre-training conditions. These ensemble voting classifiers were explored via both unweighted voting on predictions between the constituent models, as well as votes which were weighted by individual model overall accuracy over its testing set.

## 3 Results and Discussion

### 3.1 Pre-Training Results

The pre-training stage of the GCN QSAR models, utilising the 9 biomedical assay datasets, resulted in peak overall accuracy scores ranging between 74%-92% (2 s.f.) for the classification-based data (Fig. 6.a). R^2^ scores however varied more significantly between the regression-based datasets (Fig. 6.b); the most optimal R^2^ scores obtained ranged between 0.02-0.59 (2 d.p.), with the binding assay datasets resulting in significantly more optimal R^2^ scores than the CompTox toxicological assay datasets.

**Figure 6:**
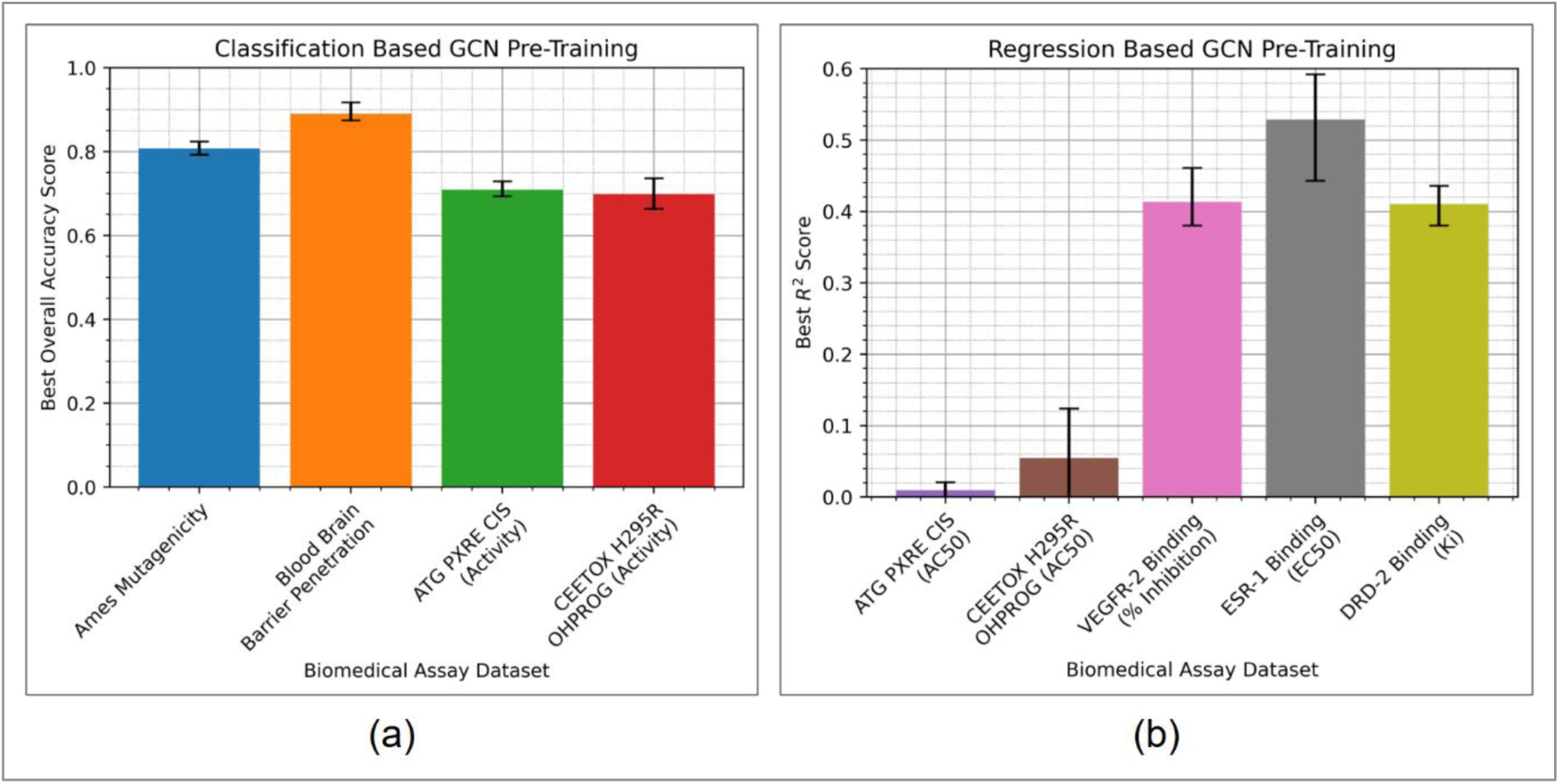
Bar charts displaying the most optimal GCN QSAR model performances, over pre-training on biomedical assay data, in terms of: (a) overall accuracy for classification-based datasets, (b) R^2^ score for regression-based datasets. Bar values are set at the average peak score for all folds, while error bars represent the range of fold-specific peak scores.

In related QSAR modelling studies, overall accuracy scores of ≥70% have typically been characterised as statistically strong for binary classification models, whereas R^2^ scores of ≥0.6 have similarly been characterised as statistically strong for regression models. There is no strict consensus on the exact performance thresholds required to deem a model as strong in its predictive capabilities, however these approximate thresholds do hold a level of consensus in literature [82-84].

Considering the above, Fig. 6.a demonstrates consistently strong GCN QSAR performances for classification tasks. While overall accuracy is often an incomplete metric for measuring predictive power of a binary classifier, due to potential imbalances between class-specific sensitivity and specificity scores, the balanced abundance of classes across folds (owing to the data stratification procedures used) mitigates this issue to a significant extent; it is highly unlikely that a sufficiently trained model, using balanced binary class training data, would result in significantly imbalanced sensitivity and specificity scores.

Fig. 6.b however demonstrates inconsistent effectiveness of the GCN QSAR model over the regression tasks; while the 3 binding assay datasets were able to result in moderately predictive models of R^2^ scores in the region of ∼0.4 to ∼0.6, the 2 CompTox toxicological assays resulted in notably poor regressor performance, with statistically insignificant R^2^ scores of <0.2. Considering the contrast in success, for the classification-based use of these same toxicological assay datasets, it may be implied that the GCN QSAR model general architecture was less appropriate for regression tasks. This is however deemed as unlikely, given the relative effectiveness of the regression model applied to the binding assay datasets. It is instead more likely that the AC50 values within the chosen CompTox assay datasets pose an inherently more challenging machine learning regression problem, compared to the binding assay datasets. Reasons for this may include inherently high-noise or erroneous experimental AC50 data, obtained through the CompTox assays, which is somewhat likely when considering the data-poor environments frequently used to estimate AC50 values from concentration-activity curves. Exact AC50 values can indeed highly vary between studies. It should also be noted that, although the regression models of the CompTox assay datasets were poor in performance, they may still potentially be of beneficial contribution in subsequent transfer learning for organ-specific toxicity, owing to potentially compatible trained functionality for processing molecular graphs and predicting toxicological outputs.

Overall, Fig. 6 indicates that the GCN QSAR model framework is effective for application over a broad range of 7 biomedical assay datasets (excluding the 2 regression-based CompTox assay datasets), which hence is an overall positive indication for subsequent use of these pre-trained model parameters for transfer learning.

### 3.2 QSAR Modelling of DILI

Numerous GCN QSAR models of DILI were obtained, via initialising different pre-training conditions prior to training on the DILI dataset (also including a lack of pre-training). For each of these DILI classifier variants, the peak overall accuracy was identified, for each fold configuration respective to each model. Comparing these scores (Fig. 7), classification performance of DILI ranges from 59%-68% (2 s.f.) overall accuracy.

**Figure 7:**
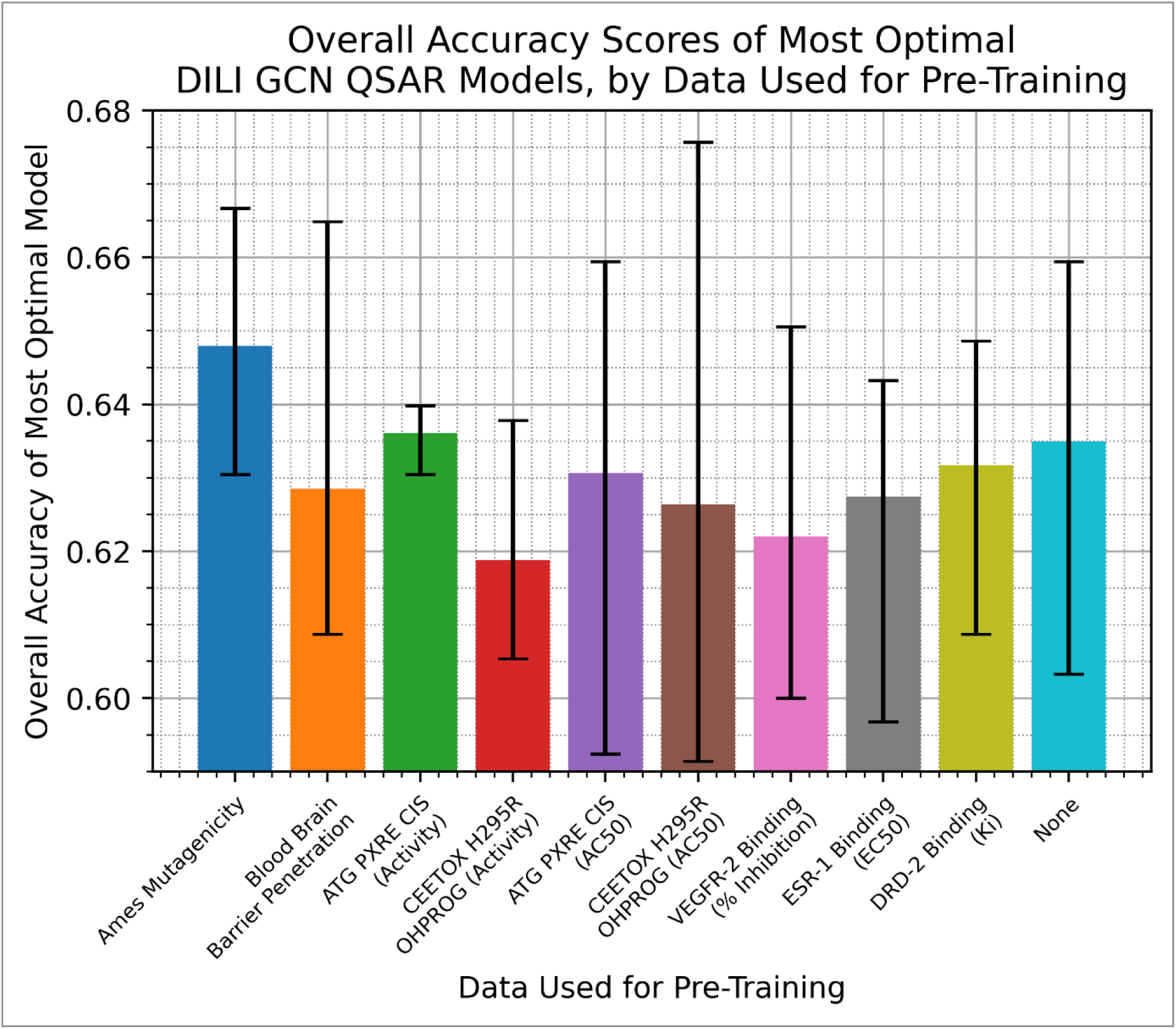
Bar chart displaying the most optimal DILI GCN QSAR model classification performances, over different pre-training conditions. Bar values are set at the average peak score for all folds, while error bars represent the range of fold-specific peak scores.

Fig. 7 demonstrates that the DILI model reached the most optimal overall accuracy of 68% (2 s.f.), when pre-trained on the CEETOX H295R OHPROG AC50 dataset. The apparent benefit of pre-training on this dataset may indicate relevance of the cytotoxicity assay to DILI, however such an explanation would be inherently *ad-hoc* in nature and neglect the similarly plausible relevance of the binary class-based assay activation dataset from the same assay, which overall underperformed for pre-training the DILI classifier. Additionally, the range of fold-specific overall accuracy scores was the widest for the model pre-trained on the CEETOX H295R OHPROG AC50 dataset, simultaneously containing the weakest scores. This hence suggests a high dependability of transfer learning model performance to specific fold-configurations, sensitive to particular classes of molecules, hence inducing biases that do not generalise uniformly well to novel data. This is apparent, when comparing to the base case of no pre-training or transfer learning; only 2 out of 9 pre-trained models matched or outperformed the base case on average overall accuracy score, while 4 out of 9 matched or outperformed when considering peak overall accuracy scores from particular fold-configurations. This indicates an overall weakly negative impact of transfer learning, on building GCN-powered QSAR models of DILI classification.

The models pre-trained on Ames mutagenicity notably outperformed the base case, via both average overall accuracy, as well as smaller variability between different fold-configurations. Although this may indicate beneficial relevance of the Ames mutagenicity endpoint to DILI, such an inference would be *ad-hoc* and negate other likely considerations involving abundance and type of molecular datapoints available in the Ames mutagenicity dataset.

### 3.3 QSAR Modelling of DIRI

Numerous GCN QSAR models of DIRI were obtained and compared, via the same approach as in Section 3.2. Comparing these scores (Fig. 8), classification performance of DIRI ranges from 65%-76% (2 s.f.) overall accuracy.

**Figure 8:**
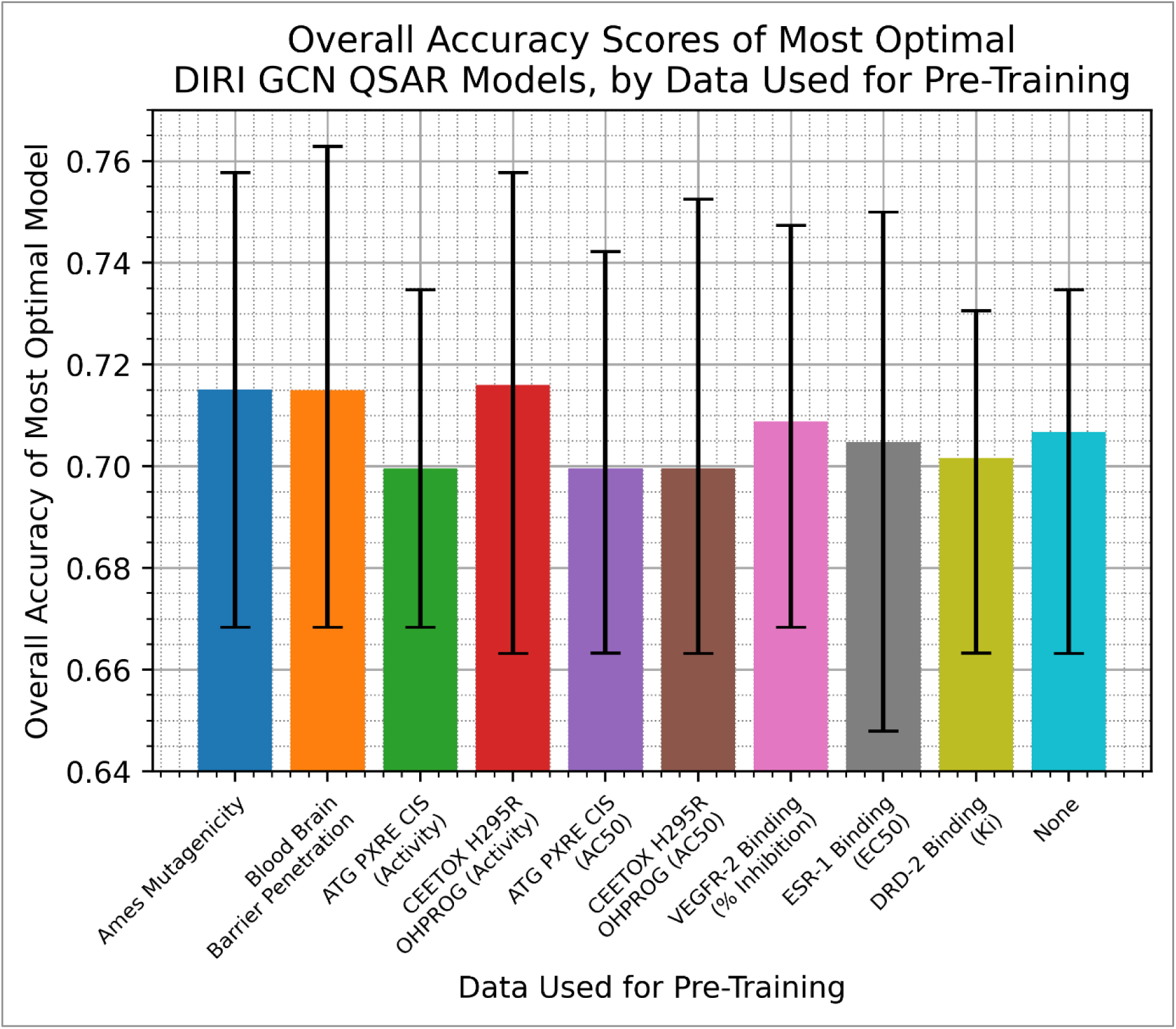
Bar chart displaying the most optimal DIRI GCN QSAR model classification performances, over different pre-training conditions. Bar values are set at the average peak score for all folds, while error bars represent the range of fold-specific peak scores.

Fig. 8 demonstrates comparably more uniform average overall accuracy scores and ranges of values, compared to Fig. 7. This may either be due to inherent differences in the modelling ease of DIRI as an endpoint, or more likely due to considerations of the DIRI dataset’s chemical space and how advantageously it may be processed by models pre-trained on the other included molecular datasets.

4 out of 9 transfer learning cases outperformed the base case, in terms of average overall accuracy score, whereas 7 out of 9 outperformed in terms of peak overall accuracy score. Fig. 8 hence contrasts the findings from Fig. 7, instead suggesting weakly positive benefit to transfer learning, at least in the context of GCN-powered QSAR modelling of DIRI classification.

The highest average overall accuracy score was 72%, obtained via transfer learning using the model pre-trained on CEETOX H295R OHPROG activation – which interestingly, is the same assay which outperformed via peak overall accuracy score in Fig. 7, albeit for the AC50 regression case. The highest peak overall accuracy score was 76%, obtained via transfer learning using the model pre-trained on classification of BBBP. This is in contrast to Fig. 7, and may suggest favourable distributions of molecular datapoints in particular fold-configurations for the BBBP dataset, which held circumstantial advantages in informing the model for handling the DIRI dataset.

Furthermore, Ames mutagenicity outperformed the base case both in terms of average and peak overall accuracy, in line with the findings of Fig. 7 (albeit to a less statistically significant extent). This may indicate that the Ames mutagenicity dataset is uniquely advantageous for transfer learning in the toxicological QSAR modelling domain, perhaps due to abundance or distribution of molecular datapoints, even if only of a statistically weak positive effect.

### 3.4 QSAR Modelling of DICT

In the same manner as in Sections 3.2 and 3.3, numerous QSAR models of DICT were obtained and compared. Comparing these scores (Fig. 9), classification performance of DICT ranges from 54%-65% (2 s.f.) overall accuracy.

**Figure 9:**
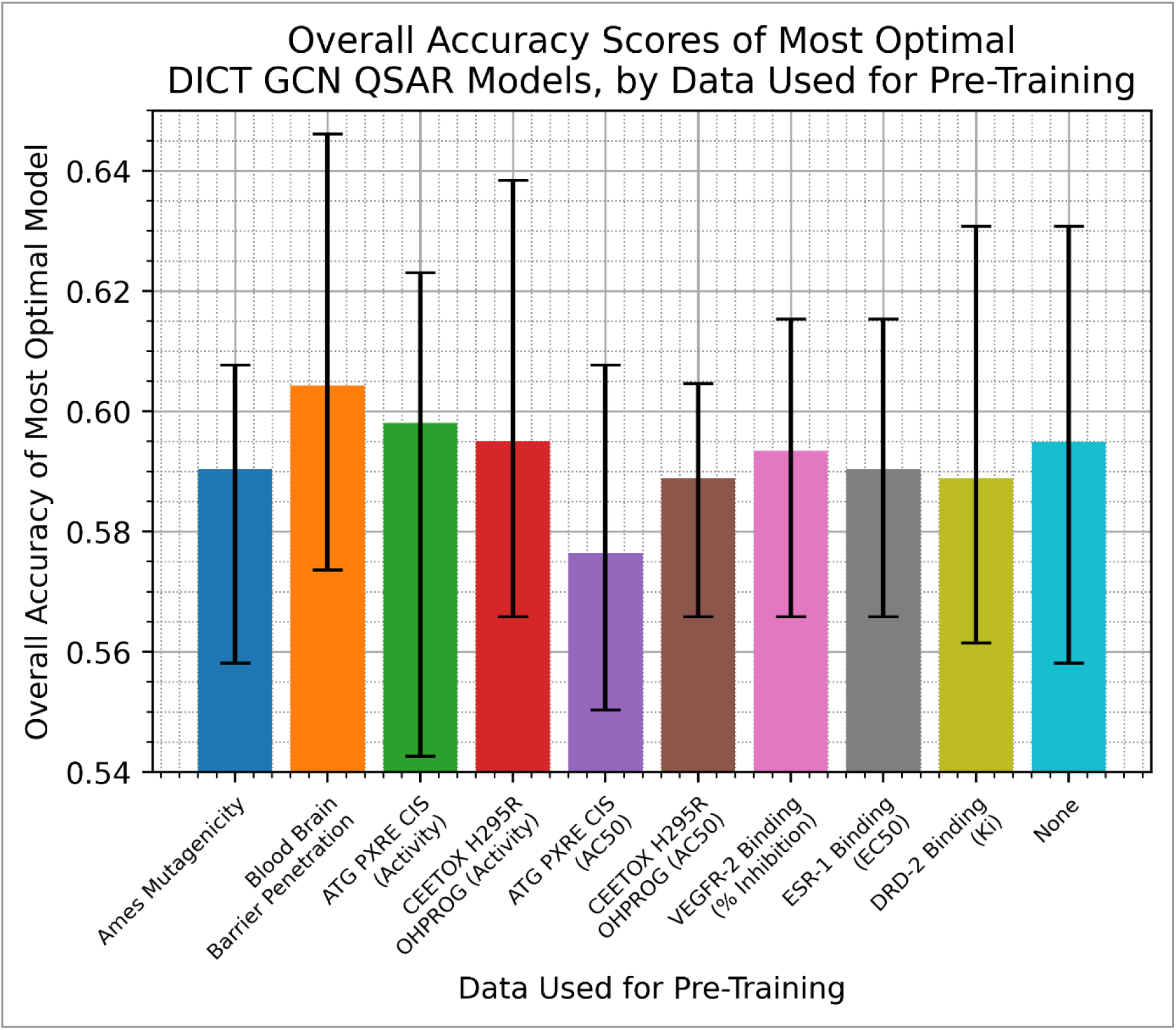
Bar chart displaying the most optimal DICT GCN QSAR model classification performances, over different pre-training conditions. Bar values are set at the average peak score for all folds, while error bars represent the range of fold-specific peak scores.

Fig. 9 demonstrates that the models pre-trained on BBBP outperformed the base case, both in terms of average overall accuracy (60%) and peak overall accuracy (65%), while furthermore resulting in higher scores across even the worst-performing fold-configurations, for both models. This aligns with the findings of Fig. 8, although contradictory of the findings of Fig. 7. In contrast to Fig. 8, however, Ames mutagenicity underperformed as a dataset for pre-training and transfer learning, across both average and peak overall accuracy metrics against the base case.

Only 3 out of 9 pre-trained models matched or outperformed the base case, in terms of average overall accuracy, whereas 3 out of 9 matched or outperformed in terms of peak overall accuracy. Interestingly, CEETOX H295R OHPROG activation was among the 3, for both metrics. Several other pre-trained cases slightly underperformed the base case, but within close statistical overlap, hence *de-facto* of the same (or otherwise closely comparable) performance. This further affirms the findings from Fig. 7, suggesting weakly negative impact of transfer learning, for creating GCN-powered QSAR models of organ-specific toxicity.

When taken in consideration with the findings of Fig. 7 and Fig. 8, an overall neutral benefit of transfer learning is uncovered, however with specific datasets (Ames mutagenicity, BBBP and CEETOX H295R OHPROG) outperforming on average. This suggests that transfer learning on GCN-powered QSAR models may be beneficial, albeit of limited statistical significance, when considering carefully chosen specific datasets that garner general benefit.

### 3.5 Applicability Domain of DILI QSAR Models

Fig. 10 displays the constructed convex hull regions for AD, as per Fig. 5, for each of the GCN QSAR models of DILI trained and tested in this study (including for different fold-configurations and pre-training conditions).

**Figure 10:**
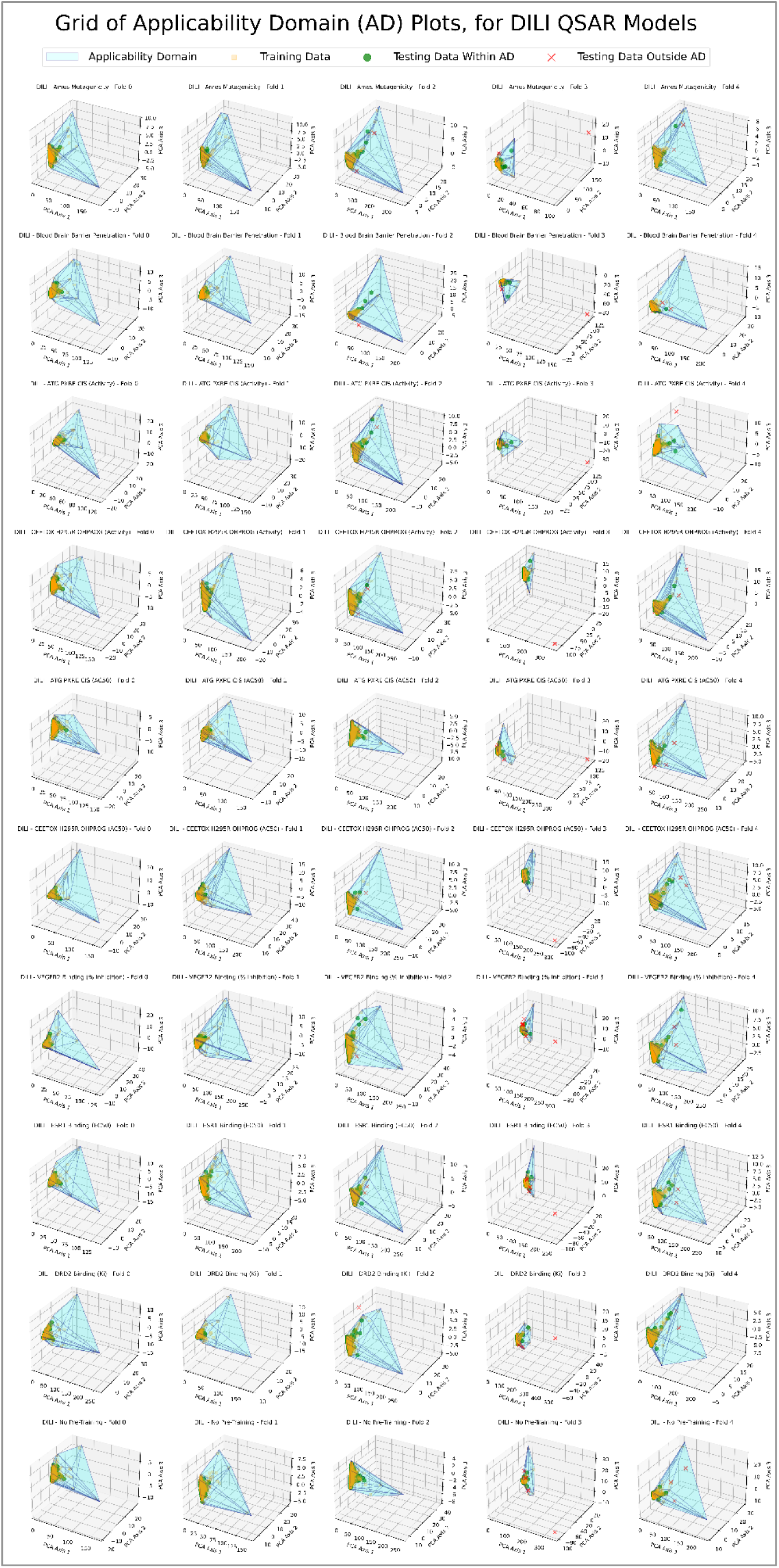
3-dimensional plots and analyses of AD regions, as per Fig. 5, for each fold-configuration of each differently pre-trained DILI GCN QSAR model.

The analyses visualised through Fig. 10 demonstrate that the majority of DILI testing data occurred within the AD of each model, hence affirming the effectiveness of the 5-fold cross validation and data stratification used, as well as the reasonable distribution of molecular datapoints in the DILI dataset. Furthermore, the reliability of the results of Fig. 7 and its subsequent analyses, are strengthened by the occurrence of the majority of testing data within the AD.

Another key finding from Fig. 10 is that similar convex hull geometric shapes were obtained (recurring fitted around dense clusters of training data embeddings, with sharp distant points from specific outliers), across same fold-configurations, for models of differing training conditions. This hence indicates that training over the DILI dataset was significantly more impactful on influencing how the models processed the testing data, compared to pre-training on differing datasets. This hence affirms previous studies that have inferred limited benefit of transfer learning for GNNs. The occurrence of certain exceptions, however, in terms of anomalous convex hull shapes which deviate significantly from other ADs for the same fold-configuration, may call into question the extent to which this holds (or alternatively, indicate a stochastic phenomenon).

### 3.6 Applicability Domain of DIRI QSAR Models

Fig. 11 presents the same analyses and visualisations as Fig. 10, but applied to the DIRI GCN QSAR models. The overall trends uncovered from Fig. 10 largely hold for Fig. 11 as well – e.g. in terms of similar convex hull shapes for the same fold configurations (even when subject to different pre-training conditions), largely affirming the notion that the transfer learning explored in this study was of limited impact. Furthermore, the majority of testing data points occurred within the evaluated AD regions.

**Figure 11:**
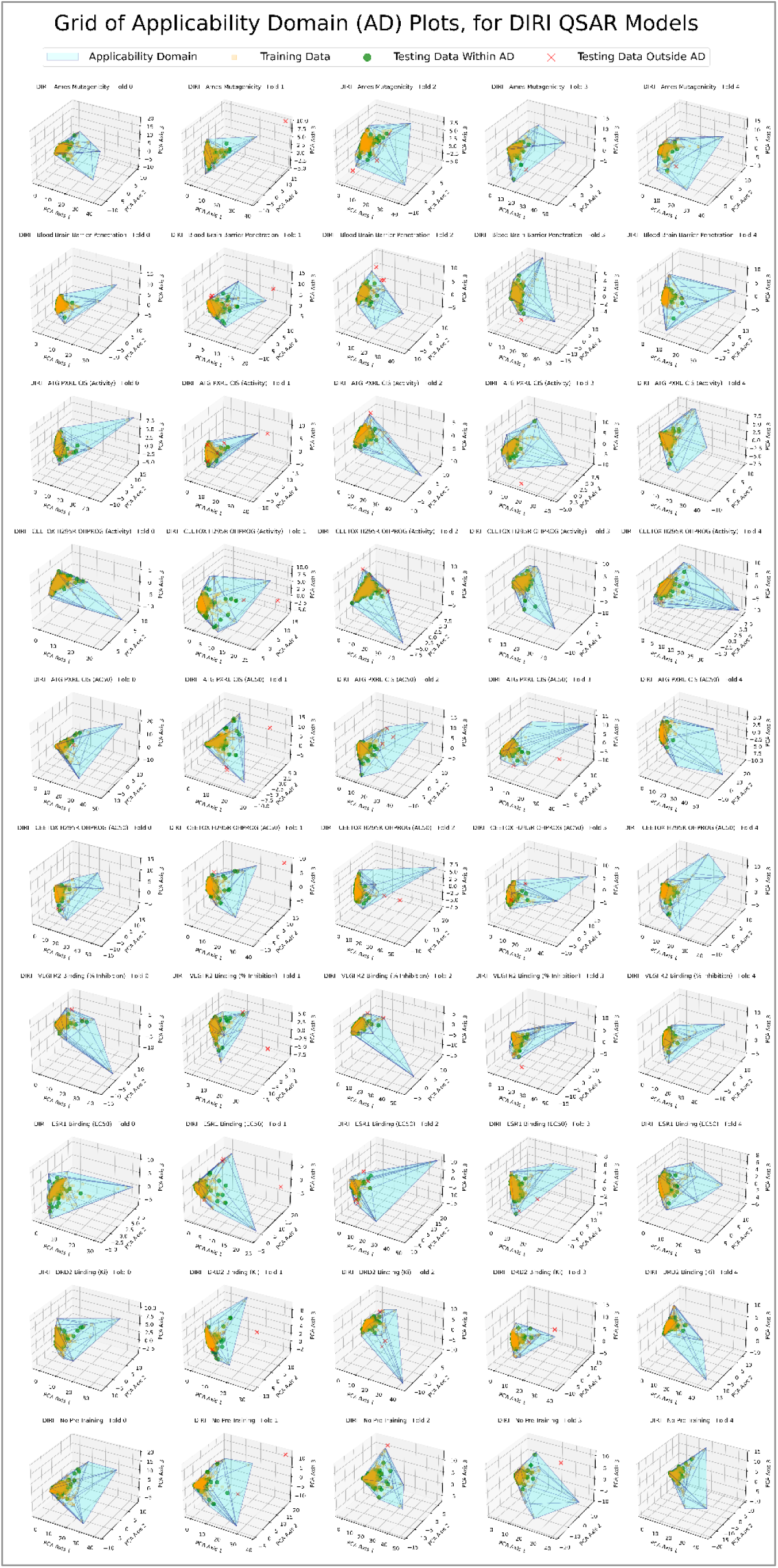
3-dimensional plots and analyses of AD regions, as per Fig. 5, for each fold-configuration of each differently pre-trained DIRI GCN QSAR model.

The recurring shapes of Fig. 11 are visibly different to those from Fig. 10, indicating that unique AD regions occurred for the different endpoint of DIRI. The convex hull shapes of Fig. 11 however follow a similar notion of largely being fit around dense clusters of embedded datapoints, with sharp vertices in the 3D convex hull geometries arising from isolated outliers. There is naturally an extent of blank space between the main bodies of the dense clusters and the isolated outliers, still considered as part of the AD – excluding outliers and tightening the AD space may hence be of benefit, for enabling more robust scrutiny of models and application to novel data.

The use of PCA, for reducing embedding dimensionality to enable 3D AD definitions, was beneficial for visualisation, but may risk loss of information and creation of overly simplistic and generous AD regions. Furthermore, distribution of embedded datapoints in the transformed 3D space is unlikely to conserve original Euclidean distances in higher dimensional latent space – but rather warp them, hence also raising validity questions about the appropriateness of using convex hulls in PCA-transformed 3D spaces. However, instead defining AD via direct calculation of convex hulls over original 200-dimensional latent spaces, would likely lead to very sharply jagged AD regions which tightly overfit to statistical noise in the conceptual hyper-surface that encapsulates the training data embeddings, likely excessively excluding novel datapoints from being able to occur within the AD. A level of de-noising (i.e. statistical simplification), via PCA (which only conserves the most consequential variance in the data), was hence likely beneficial in creating more pragmatic and useful AD definitions.

### 3.7 Applicability Domain of DICT QSAR Models

As per the visualisations of the AD for Fig. 12, comparably findings apply to the DICT GCN QSAR models as those for DILI (Fig. 10) and DIRI (Fig. 11) – i.e. similar convex hull shapes over particular fold configurations, largely irrespective of different pre-training conditions, as well as the majority of testing data having occurred within the AD of each model.

**Figure 12:**
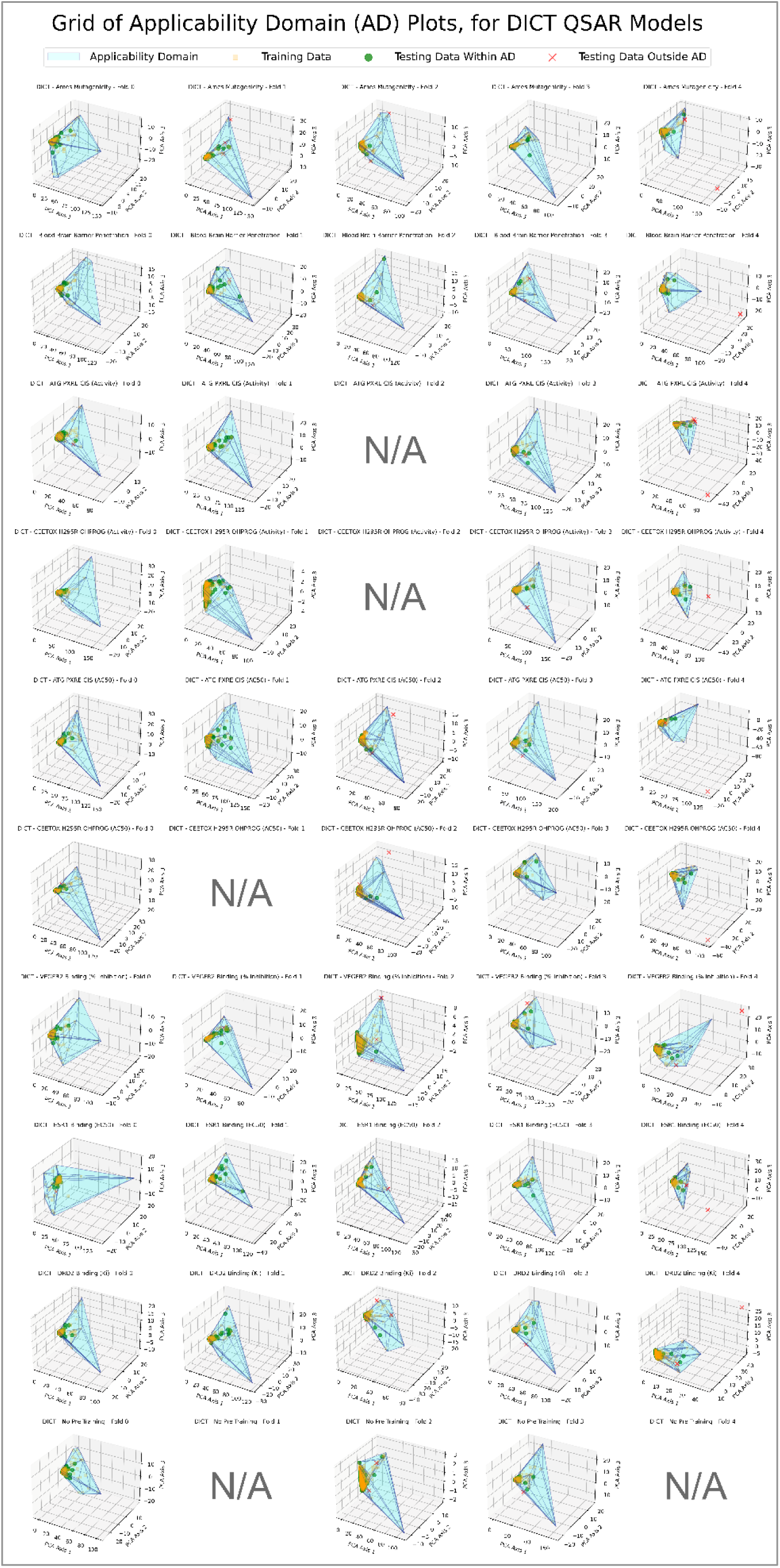
3-dimensional plots and analyses of AD regions, as per Fig. 5, for each fold-configuration of each differently pre-trained DICT GCN QSAR model.

Despite these initial findings, supporting the notion of transfer learning having had only minimal impacts on final DICT GCN QSAR model trained states, certain AD plots are marked as “N/A” in Fig. 12 – due to no model improvement having been reached (in terms of overall accuracy over the testing set, over any of the training epochs). This hence indicates that the DICT models were more problematic than the DILI or DIRI models, in training and testing, possibly owing to the DICT dataset being less easy to model (which is supported by the lower peak and mean overall accuracy scores reached). While the DICT dataset may have been inherently more challenging for GCN QSARs to model, it is also an inherently smaller dataset than the DILI or DIRI datasets used – hence potential posing the limitations of a few-shot learning AI modelling task, when used on conventional architectures. Our previous research has indicated that GCNs are not an optimal GNN architecture for data-scarce toxicological data environments, compared to other architectures such as Graph Attention Networks (GATs) and Graph Isomorphism Networks (GINs).

Furthermore, contrary to the findings from Fig. 10 and 11, Fig. 12 does demonstrate some notable deviations in AD convex hull shapes, for the same fold configurations – particularly for folds 0 and 4. This may further indicate a lack of suitable model optimisation, over the training epochs (even if some optimisation of overall accuracy over the testing set did occur), hence affirming that the DICT models were trained less effectively than the DILI and DIRI models.

### 3.8 Predictions for SARMs

The most optimal models (in terms of test overall accuracy score) were applied to predict for 25 SARMs – the results of which are displayed in Table 1. The most optimal DILI model was pre-trained on the CEETOX_H295R_OHPROG assay (using AC50 values), using fold configuration 0. For DIRI, the selected model was the one pre-trained on BBBP, using fold configuration 4, whereas for DICT the selected model was also pre-trained on BBBP, but using fold configuration 0 instead.

**Table 1:**
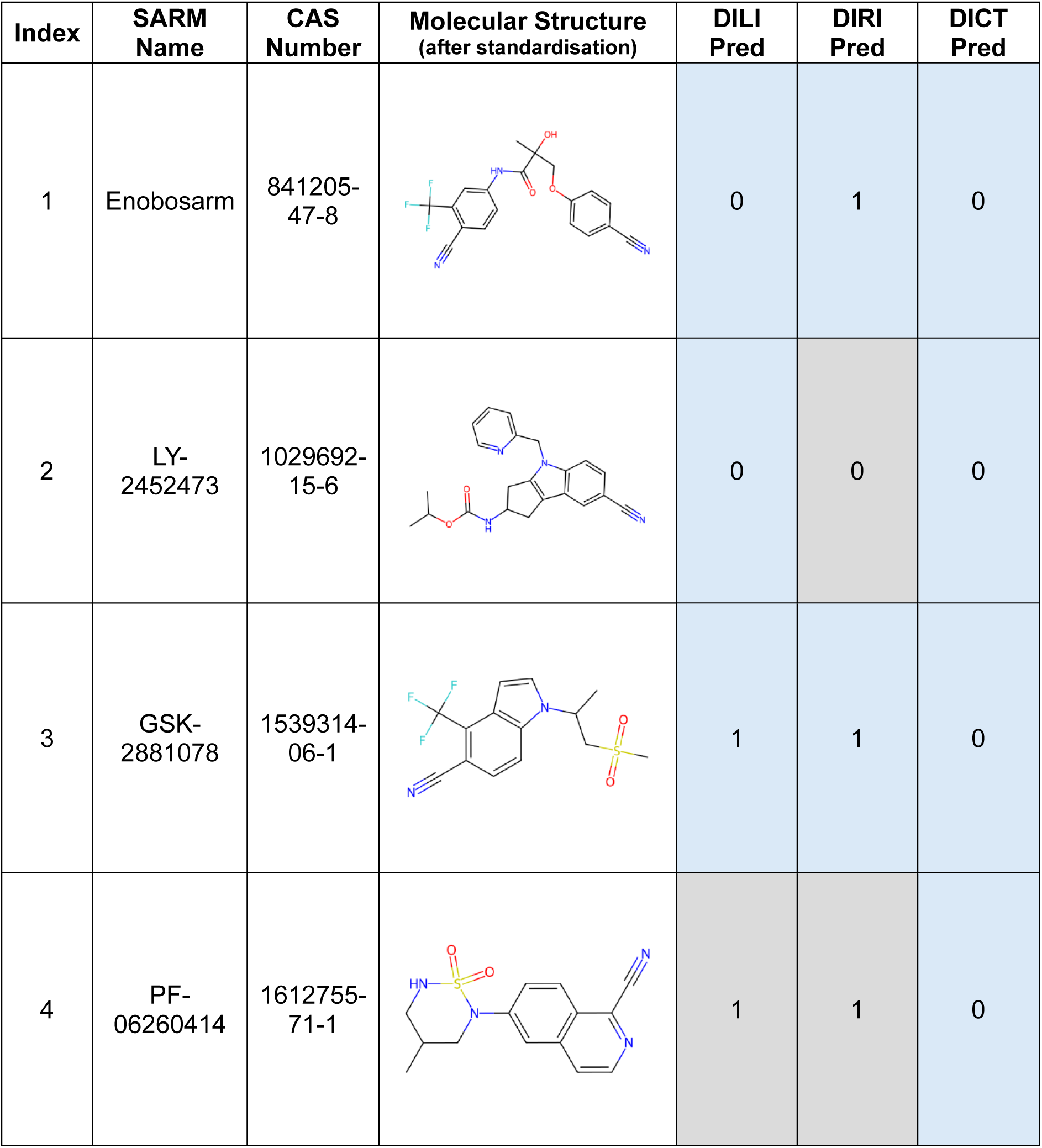
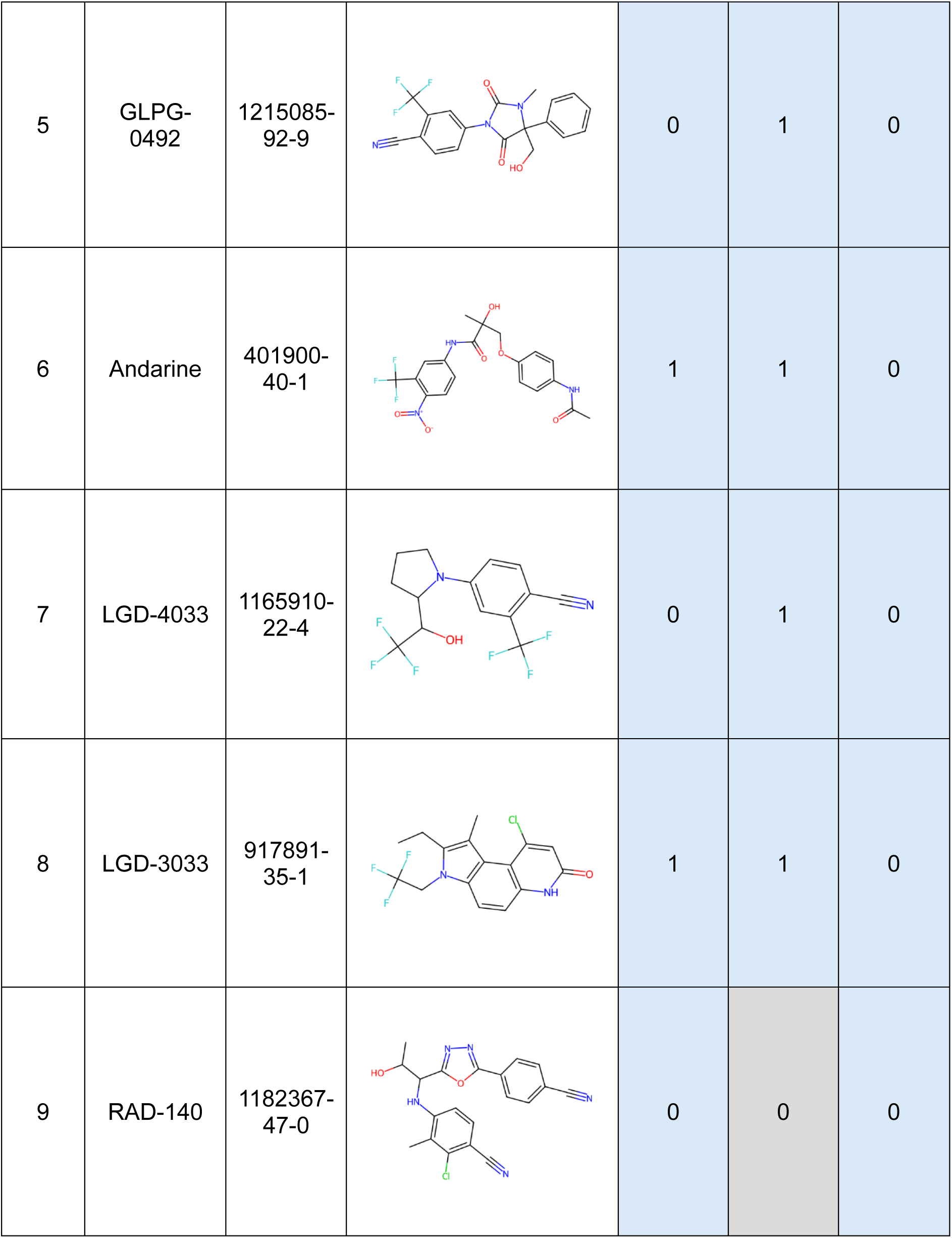
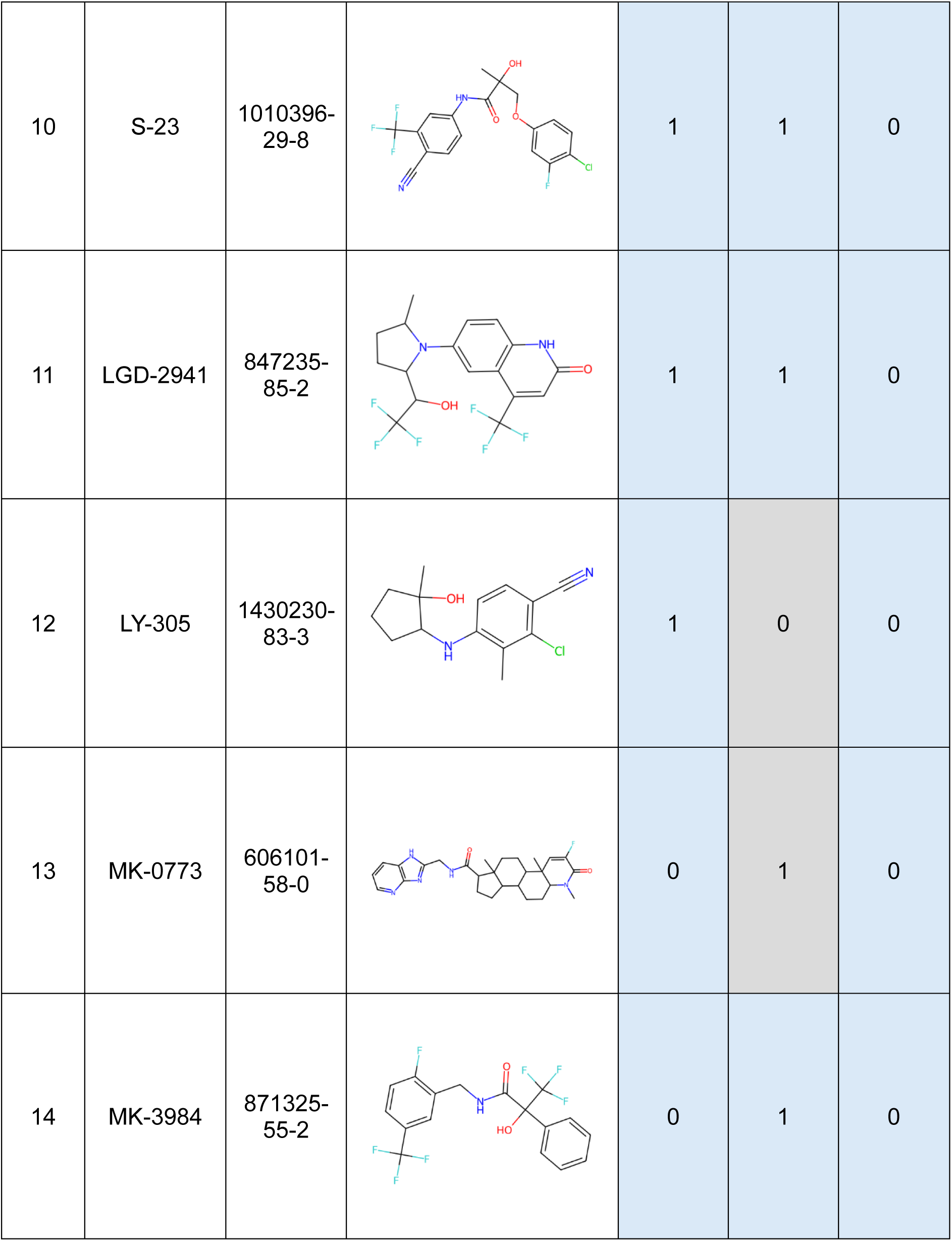
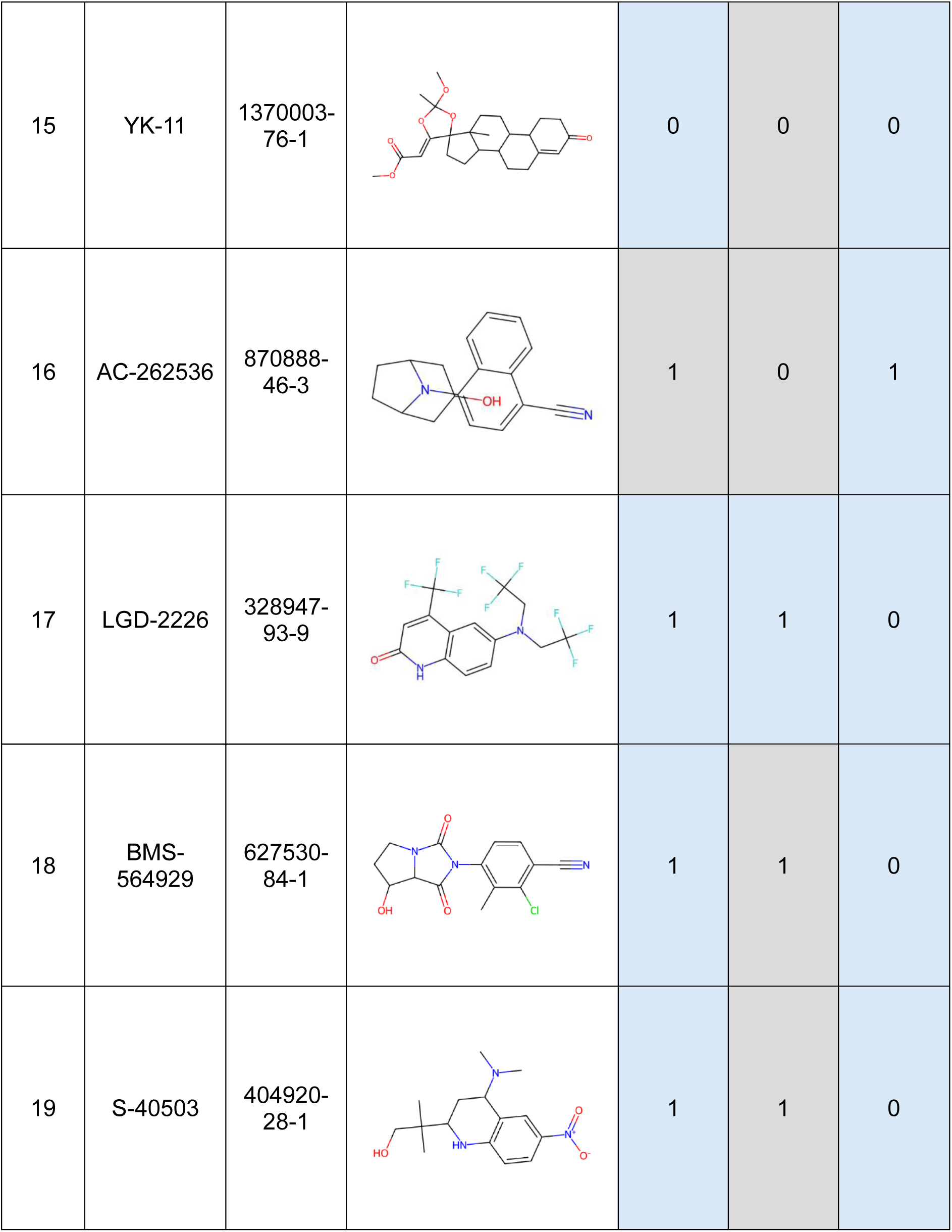
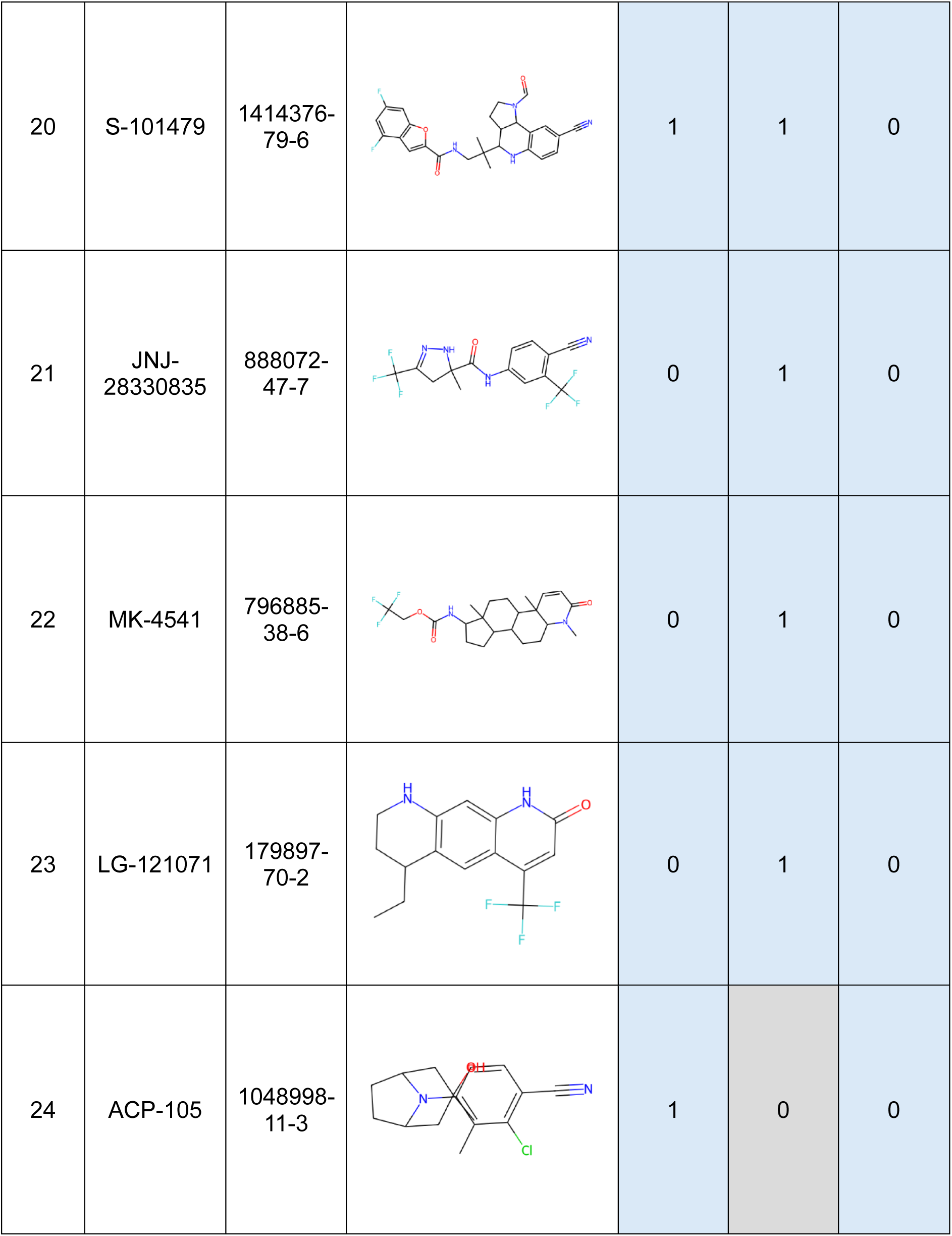
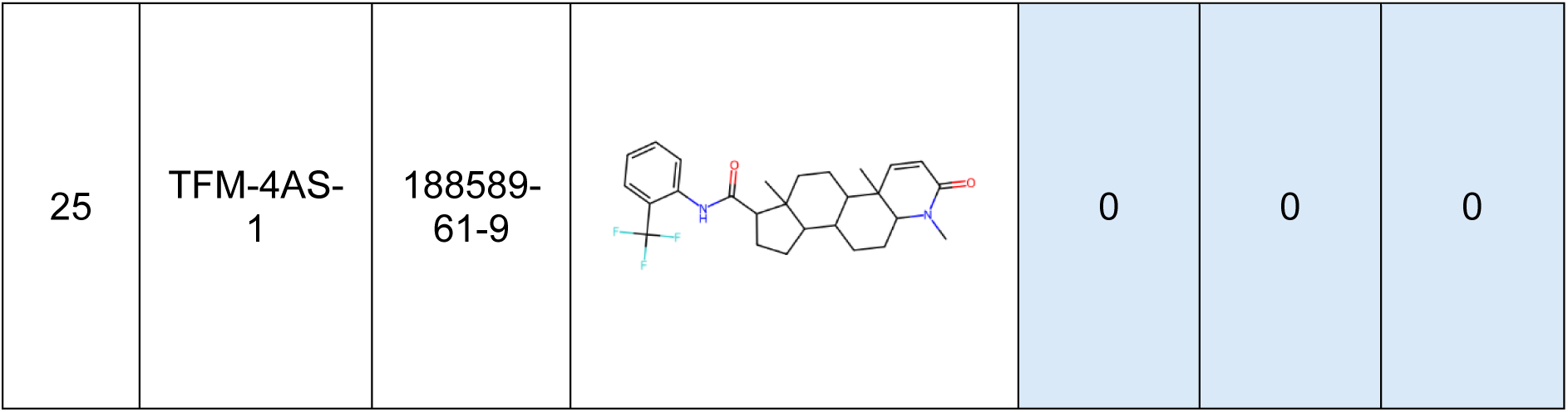
Details of the SARMs investigated in this study, along with corresponding DILI, DIRI and DICT predictions, by the highest overall accuracy models from Fig. 7-9. Predictions that were within the AD, are highlight in pale blue, whereas outside cases are highlighted grey.

| Index | SARM Name | CAS Number | Molecular Structure (after standardisation) | DILI Pred | DIRI Pred | DICT Pred |
| --- | --- | --- | --- | --- | --- | --- |
| 1 | Enobosarm | 841205-47-8 |  | 0 | 1 | 0 |
| 2 | LY-2452473 | 1029692-15-6 |  | 0 | 0 | 0 |
| 3 | GSK-2881078 | 1539314-06-1 |  | 1 | 1 | 0 |
| 4 | PF-06260414 | 1612755-71-1 |  | 1 | 1 | 0 |
| 5 | GLPG-0492 | 1215085-92-9 |  | 0 | 1 | 0 |
| 6 | Andarine | 401900-40-1 |  | 1 | 1 | 0 |
| 7 | LGD-4033 | 1165910-22-4 |  | 0 | 1 | 0 |
| 8 | LGD-3033 | 917891-35-1 |  | 1 | 1 | 0 |
| 9 | RAD-140 | 1182367-47-0 |  | 0 | 0 | 0 |
| 10 | S-23 | 1010396-29-8 |  | 1 | 1 | 0 |
| 11 | LGD-2941 | 847235-85-2 |  | 1 | 1 | 0 |
| 12 | LY-305 | 1430230-83-3 |  | 1 | 0 | 0 |
| 13 | MK-0773 | 606101-58-0 |  | 0 | 1 | 0 |
| 14 | MK-3984 | 871325-55-2 |  | 0 | 1 | 0 |
| 15 | YK-11 | 1370003-76-1 |  | 0 | 0 | 0 |
| 16 | AC-262536 | 870888-46-3 |  | 1 | 0 | 1 |
| 17 | LGD-2226 | 328947-93-9 |  | 1 | 1 | 0 |
| 18 | BMS-564929 | 627530-84-1 |  | 1 | 1 | 0 |
| 19 | S-40503 | 404920-28-1 |  | 1 | 1 | 0 |
| 20 | S-101479 | 1414376-79-6 |  | 1 | 1 | 0 |
| 21 | JNJ-28330835 | 888072-47-7 |  | 0 | 1 | 0 |
| 22 | MK-4541 | 796885-38-6 |  | 0 | 1 | 0 |
| 23 | LG-121071 | 179897-70-2 |  | 0 | 1 | 0 |
| 24 | ACP-105 | 1048998-11-3 |  | 1 | 0 | 0 |
| 25 | TFM-4AS-1 | 188589-61-9 |  | 0 | 0 | 0 |

The results of Table 1 are more concisely visualised in the Venn diagram of Fig. 13.

**Figure 13:**
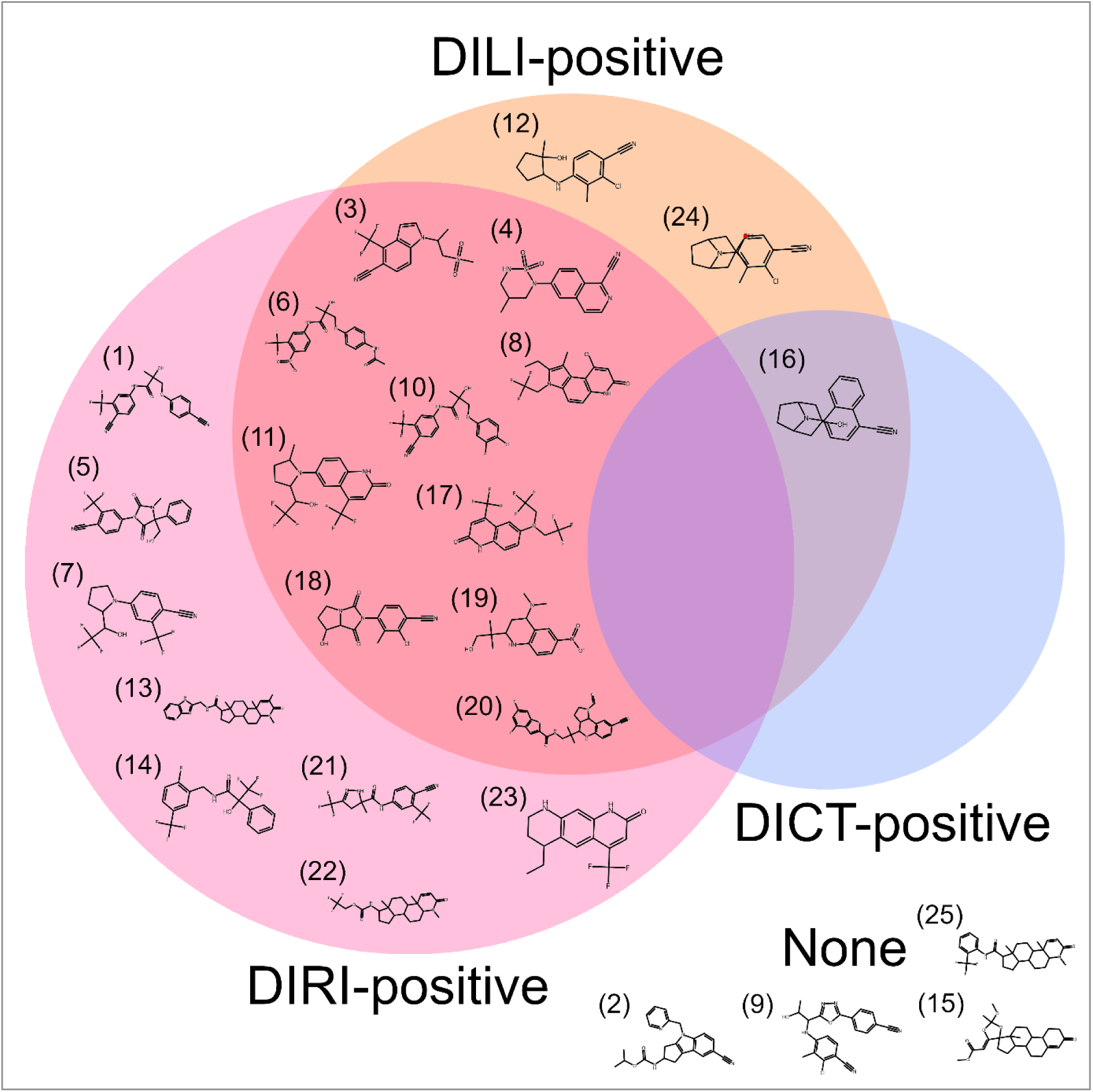
Venn diagram concisely visualising the predicted results from Table 1 (DILI, DIRI and DICT predictions for 25 SARMs). For each displayed molecular graph, the corresponding index from Table 1 is denoted at the upper left.

The results from Table 1 and Fig. 13 reveal that a majority of SARMs, considered in this study, were predicted as DIRI-positive (18 out of 25 SARMs – hence 72% of the SARMs). This raises concern, given the rising use of SARMs in sports doping. Furthermore, 13 out of 25 of the SARMs (i.e. 52% - a majority) were predicted as DILI-positive, further raising concern. Only 1 SARM was predicted as DICT-positive (while simultaneously predicted as DILI-positive), hence suggesting that DICT is not an endpoint as relevant in concern as DIRI and DILI.

Overall, only 4 out of 25 SARMs were not predicted as toxic over any of the organ-specific endpoints – hence the overwhelming majority (21 out of 25 SARMs – i.e. 84%) are either DIRI-positive, DILI-positive or both. Exactly 10 out of 25 (40%) were predicted as simultaneously DIRI-positive and DILI-positive.

Nephrotoxicity (via DIRI) is hence identified by this study as the most concerning toxicological endpoints for SARMs, as a class of chemicals, followed by hepatotoxicity (via DILI). Cardiotoxicity (via DICT) is however deemed by this study as unlikely to be a particularly concerning endpoint, for users of SARMs.

Adding further consensus of reliability to these predictions, the majority of SARMs were situated within the AD for the optimal DILI, DIRI and DICT models used (with specific locations within the embedding space visualised in Fig. 14 below).

**Figure 14:**
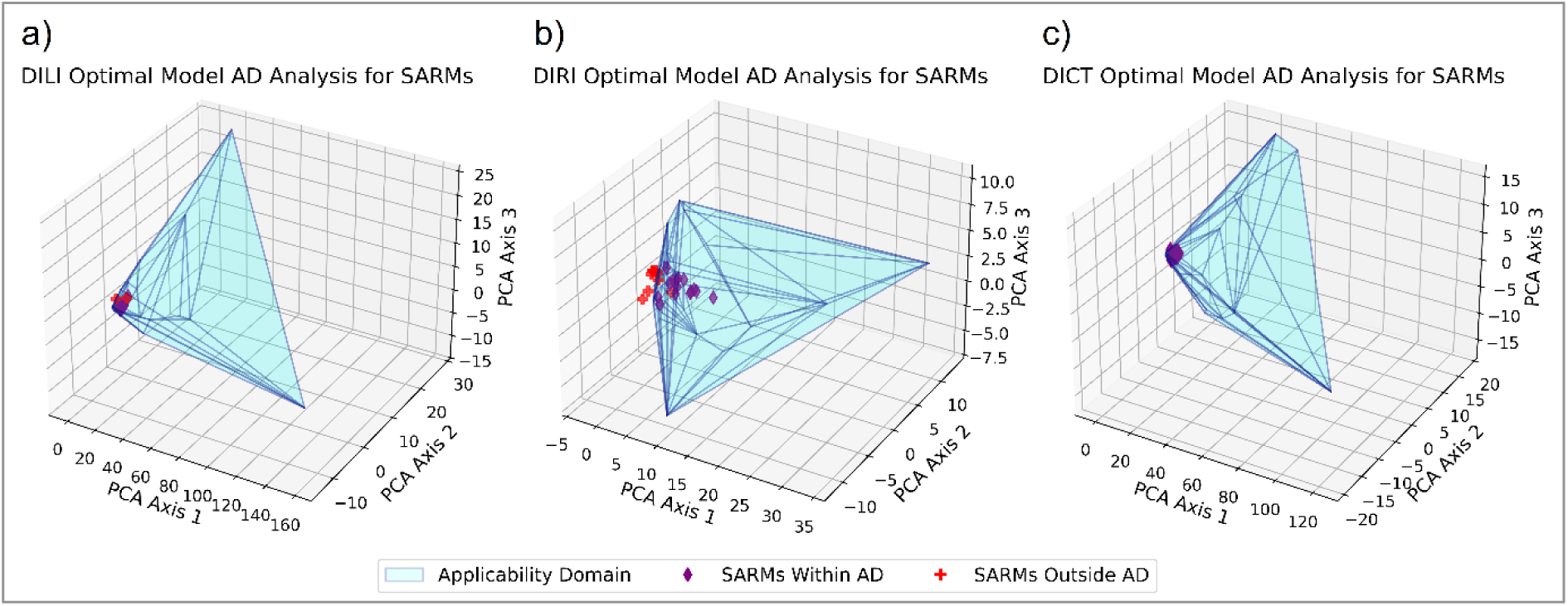
AD Plots for the most optimal single QSAR models of: a) DILI; b) DIRI; c) DICT - with the positions of the externally evaluated SARMs visualised.

The DIRI model however resulted in a sizable minority of SARMs not occurring within the AD, hence reducing the certainty that may be assigned to affected predictions (albeit nonetheless with a clear consensus of 14 SARMs, predicted as DIRI-positive, occurring within the DILI GCN QSAR model AD and hence posing as credible predictions).

Fig. 14 however may call into question the suitability of the AD definition used, given that the majority of SARMs outside of the DIRI QSAR AD (as well as the 2 SARMs outside of the DILI QSAR AD) closely clustered around a central region of the embedding space also apparent in Fig. 10, Fig. 11 and Fig. 12., in comparatively close proximity to other molecules within the AD, whereas the AD region itself stretches over blank space, to capture outliers within the training sets. A more robust definition of the AD may hence entail a density-based definition of the borders, rather than simply encapsulating all training data points with a convex hull, as well as greater statistical lenience for nearby data points (rather than the ±10% used in this study’s definition of the AD).

### 3.9 Results of Explored Ensemble Models

Fig. 15 demonstrates no clear dominance of either of the two explored EVC approaches (equal voting and weighted voting), for predicting DILI, against the overall accuracy scores of the best single model. A marginally higher peak overall accuracy score (still rounding to 68%) was however obtained via use of the weighted EVC over fold configuration 0.

**Figure 15:**
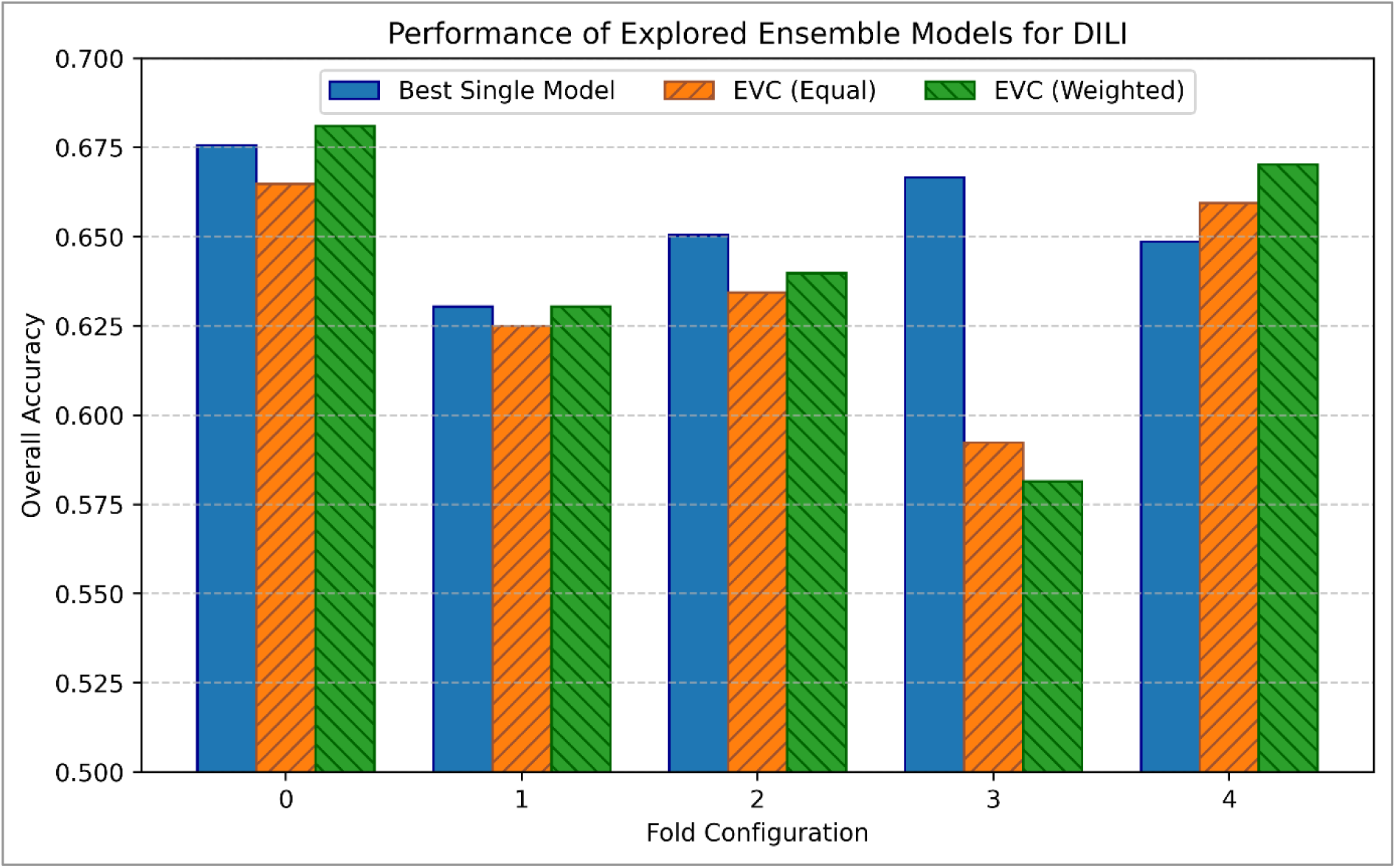
Bar chart displaying overall accuracy scores of explored EVCs, using DILI models trained in this study.

Fig. 16, similarly to Fig. 15, indicates no clear dominance of either of the two explored EVC approaches, for predicting DIRI, against the performance of the best single model. Peak overall accuracy was however mildly improved via both EVCs over fold configuration 4, reaching 77% (2 s.f.).

**Figure 16:**
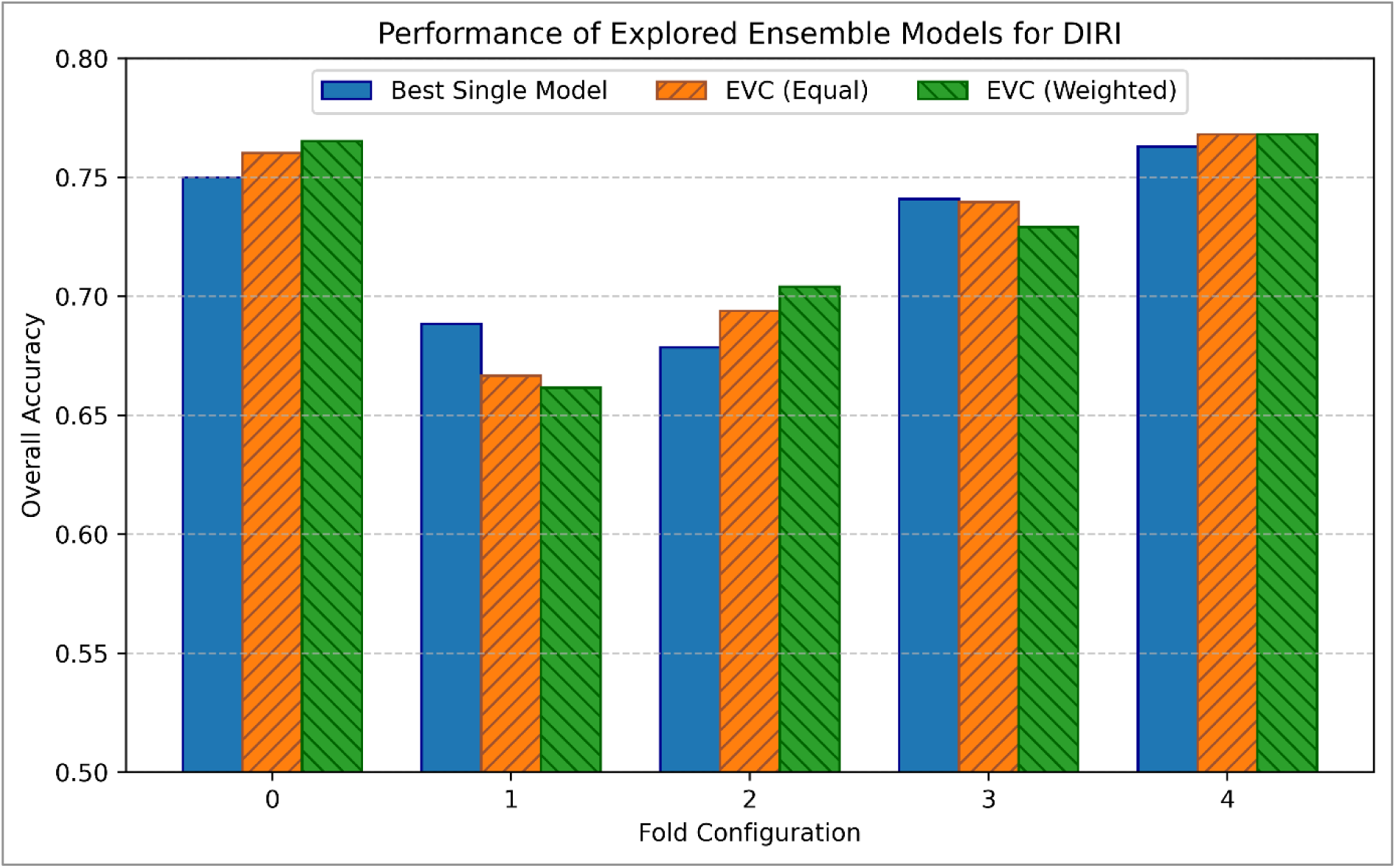
Bar chart displaying overall accuracy scores of explored EVCs, using DIRI models trained in this study.

Fig. 17, similarly to Fig. 15 and Fig. 16, fails to indicate consistent advantages of either of the two explored EVC approaches, for predicting DICT, against the performance of the best single model – although peak overall accuracy (over fold configuration 0) was significantly improved (compared to the incremental improvements visible in Fig. 15 and Fig. 16). Peak overall accuracy was improved to 67% (2 s.f.) – a 2% increase from the best single model score of 65%.

**Figure 17:**
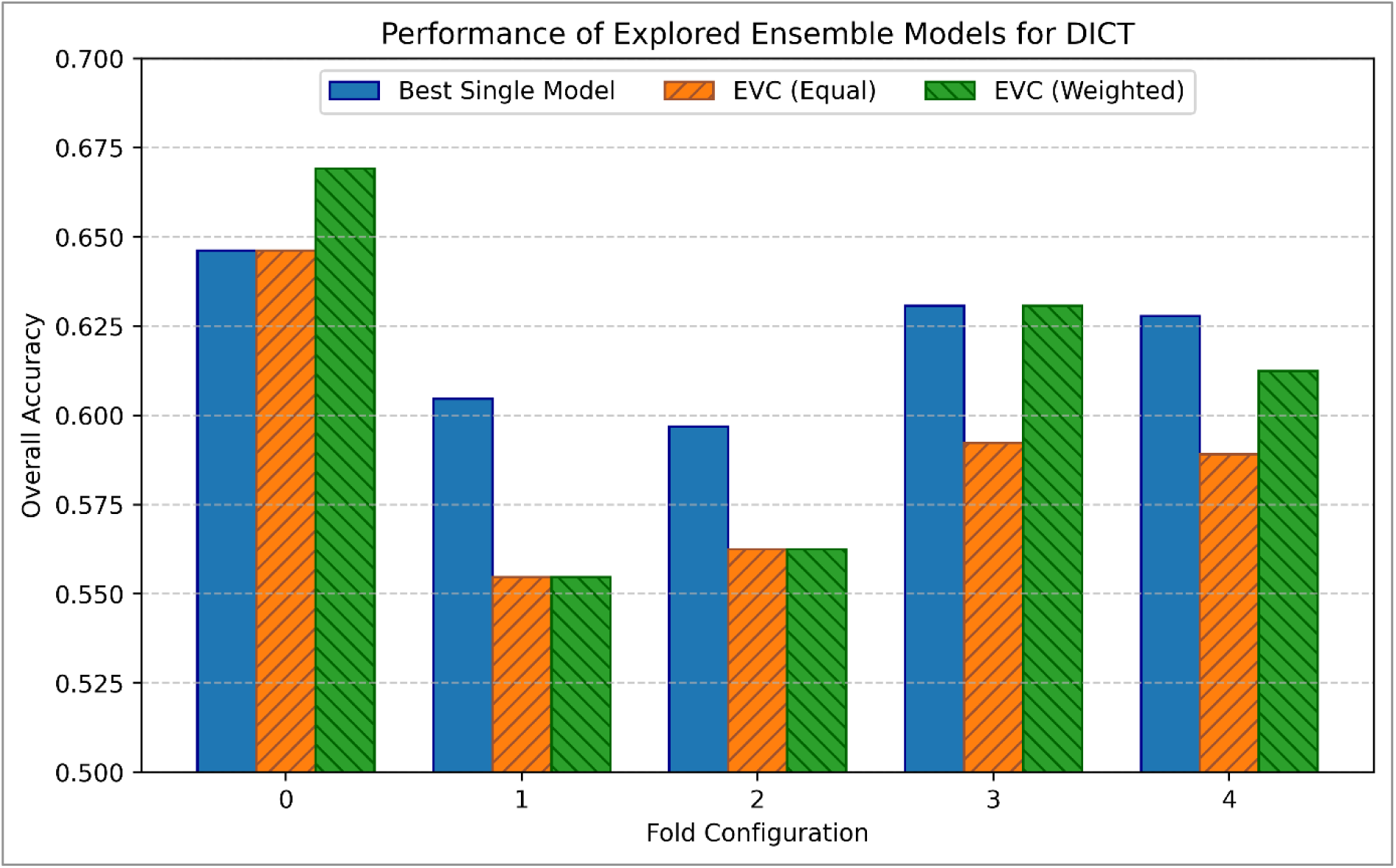
Bar chart displaying overall accuracy scores of explored EVCs, using DICT models trained in this study.

Although more significant in performance (as well as for an endpoint for which the methodology of this study otherwise underperformed, comparatively to DILI and DIRI, hence potentially gaining more notable upside from the potential use of EVCs), the weighted EVCs were not applied to predict DICT (or any other endpoint) for the SARMs, due to potential bias issues stemming from gauging testing accuracy, via voting in ensembles weighted by testing accuracy scores priorly known. This phase of the study should hence be deemed as exploratory only, providing superficial insights into the potential benefits of using ensemble modelling, incorporating voting consensuses from otherwise identical models which were subjected to different pre-training conditions.

This phase of the study however finds limited plausibility of EVCs posing any statistically significant advantage, over constituent models that were subject to different pre-training conditions. This hence indicates either that the pre-training insights were overwritten by subsequent training, or were not relevant or effective for informing the models with diversified biochemical pattern recognition capabilities.

## 4 Conclusions

From our various results and analyses, it may be concluded that SARMs pose a significant human health concern, especially for nephrotoxicity, as well as hepatotoxicity to a more moderate extent; the vast majority of SARMs (21 out of 25) were predicted as either DIRI-positive, DILI-positive, or both. Conversely, SARMs are less likely to pose a significant hazard in terms of cardiotoxicity, although one SARM was predicted as DICT-positive. Future research would benefit from further investigation of these SARMs for nephrotoxicity and hepatotoxicity, especially in more conclusive *in-vitro* and *in-vivo* studies. Future *in-silico* studies may also pose as beneficial to further investigating the potential nephrotoxicity and hepatotoxicity of SARMs, via use of more mechanistically interpretable models, as well as investigations of toxicokinetic behaviours.

The GCN-driven models of DILI, DIRI and DICT obtained were demonstrably effective, significantly outperforming the predictive power of a random guess (expectedly 50%, on a balanced binary classification problem), performing at overall accuracy scores of 76% for DIRI (improved to 77% via ensemble modelling) and 65% for DICT (improved to 67% via ensemble modelling). Further improvements to these performances may be potentially obtained via hyperparameter optimisation (perhaps through Bayesian optimisations of hyperparameters such as number and size of layers, learning rate, batch size etc.), as well as exploration of more advanced and expressive GNN algorithms, such as graph attention networks, graph isomorphism networks, graph transformer networks etc.

The novel AD definitions used, while uniquely relevant to these models, reproducible and intuitively visualisable, however held limitations in terms of physicochemical interpretability, as well as extensive low-density regions due to strict inclusion of even outlier molecules that occurred in the training set. Future expansions of this study therefore may benefit from incorporation of a more density-based paradigm for defining the AD.

The exploratory use of transfer learning on GCN-based QSAR models, to harness beneficial insights into molecular graph pattern recognition and property prediction, from randomised biomedical datasets, was largely a null result; no significant benefits were obtained. While certain pre-trained models outperformed the base case (devoid of prior pre-training), these instances appeared to demonstrate only marginal improvements in overall accuracy, as well as inconsistently across different fold configurations and toxicological endpoints. This affirms existing research in the wider topic of transfer learning on GCNs, outlining the limited benefits which are potentially fundamental to the GCN architecture itself. A similar null result was also obtained for exploration of EVCs, with only marginal and inconsistent improvements obtained, further affirming the notion of different instances of transfer learning on GCNs having offered statistically negligible benefits, if any.

Future expansions of the study may hence also benefit from exploring the possibility of more effective transfer learning for toxicological QSAR models which use more computationally intensive AI architectures that are more likely to be improved by such a methodology, such as language-based transformer models, or more expressive graph neural networks which also incorporate 3-dimensional coordinate information (such as equivariant graph neural networks) and even quantum-mechanical information.

## Declarations

### Availability of Data and Materials

All software developed, as well as data and other materials, are available in the following GitHub folder (constituent of the wider TOX-AI GitHub repository): https://github.com/alexanderkalian/TOX-AI/tree/main/transfer_learning_gcns

### Competing Interests

The authors of this study declare no competing interests.

### Funding

This work was supported by grants from the Biotechnology and Biological Sciences Research Council [grant number BB/T008709/1] and the Food Standards Agency [Agency Project FS900120].

### Authors’ Contributions

ADK and ACS conceptualised the study. CH, MG, OJO, EB, CP and JLCMD supervised the study. ACS and JL provided non-supervisory advice and guidance, for the study. ADK carried out computational experiments for the study. ADK produced analyses of the results. ADK, ACS and OJO contributed to writing the manuscript. ADK produced diagrams for the manuscript. ADK, ACS, JL, OJO, MG and CH reviewed and edited the manuscript.

## Acknowledgements

The content of this article does not necessarily reflect the views of the Food Standards Agency.

The views expressed in this article do not reflect the views of the European Food Safety Authority (EFSA) and/or are a reflection of the views of the authors only.

This paper aims to contribute to the international network on Advancing the Pace of Chemical Risk Assessment (APCRA), to contribute to the use of New Approach Methodologies (NAMs) in chemical risk assessment and ultimately reduce animal testing.

## Notes

https://github.com/alexanderkalian/TOX-AI/tree/main/transfer_learning_gcns

### Competing Interest Statement

The authors have declared no competing interest.

